# Distinguishing the signatures of local environmental filtering and regional trait range limits in the study of trait-environment relationships

**DOI:** 10.1101/528836

**Authors:** Pierre Denelle, Cyrille Violle, François Munoz

**Affiliations:** CEFE UMR 5175, CNRS - Université de Montpellier - Université Paul-Valéry Montpellier – EPHE -1919 route de Mende, F-34293 Montpellier, CEDEX 5, France; University Grenoble-Alpes, LECA, 2233 Rue de la Piscine, 38041 Grenoble Cedex 9, France

**Keywords:** community assembly, functional biogeography, environmental filtering

## Abstract

Understanding the imprint of environmental filtering on community assembly along environmental gradients is a key objective of trait-gradient analyses. Depending on local constraints, this filtering generally entails that species departing from an optimum trait value have lower abundances in the community. The Community-Weighted Mean (CWM) and Variance (CWV) of trait values are then expected to depict the optimum and intensity of filtering, respectively. However, the trait distribution within the regional species pool and its limits can also affect local CWM and CWV values apart from the effect of environmental filtering. The regional trait range limits are more likely to be reached in communities at the extremes of environmental gradients. Analogous to the mid-domain effect in biogeography, decreasing CWV values in extreme environments can then represent the influence of regional trait range limits rather than stronger filtering in the local environment. We name this effect the “Trait-Gradient Boundary Effect” (TGBE). First, we use a community assembly framework to build simulated communities along a gradient from a species pool and environmental filtering with either constant or varying intensity while accounting for immigration processes. We demonstrate the significant influence of TGBE, in parallel to environmental filtering, on CWM and CWV at the extremes of the environmental gradient. We provide a statistical tool based on Approximate Bayesian Computation to decipher the respective influence of local environmental filtering and regional trait range limits. Second, as a case study, we reanalyze the functional composition of alpine plant communities distributed along a gradient of snow cover duration. We show that leaf trait convergence found in communities at the extremes of the gradient reflect an influence of trait range limits rather than stronger environmental filtering. These findings challenge correlative trait-environment relationships and call for more explicitly identifying the mechanisms responsible of trait convergence/divergence along environmental gradients.

## Introduction

Quantifying the physiological responses of organisms and communities along environmental gradients is pivotal in ecology and biogeography (Lomolino et al. 2006, Violle et al. 2014). However, we know little about the sensitivity of such responses to environmental or physiological limits, i.e. to boundary effects. Boundary effects have been broadly addressed in biogeography, in terms of taxonomic diversity at the limits of environmental gradients. Specifically, the mid-domain effect represents an artefactual peak of species richness at the center of latitudinal gradient (Colwell and Lees 2000, Colwell et al. 2004) or of species range at the center of an environmental gradient (Letten et al. 2013) due to sampling issues. Here we recast this hypothesis through the lens of trait-based ecology. More specifically, we argue that the parameters of the local trait distribution at the edge of environmental and/or trait gradients can be misinterpreted because the regional trait distribution is not properly quantified. While the influence of the taxonomic composition and richness of a source species pool on local community assembly have received much interest, the influence of the functional composition of the pool has only recently come to focus (Patrick and Brown 2018, Spasojevic et al. 2018). This influence, that we coined ‘trait-gradient boundary effect’ (TGBE), can combine with the effect of environmental filtering, as both constrain the moments of the local trait distribution at community scale (Kraft et al. 2015). We here provide a method to separate the influence of environmental filtering on local community assembly from the imprint of regional trait distribution, in order to avoid misinterpretations on the strength of environmental filtering.

Functional traits are attributes reflecting the ability of individuals to survive and reproduce in a local environment (Violle et al. 2007). Assembly processes shape the distribution of functional trait values within communities (McGill et al. 2006), and in particular environmental filtering represents the control of the local trait distribution by abiotic factors (Kraft et al. 2015). Environmental filtering generally includes two components (Shipley 2010): (i) an optimal trait value or combination allowing maximal performance and greater abundance in the community, and (ii) an intensity value quantifying how sharp the decrease of species performance around the optimal trait value is (Fig. 1). Varying the functional composition of communities along environmental gradients is then expected to reflect changing optimal values and/or filtering intensity (Ackerly and Cornwell 2007). Because the variation of performance around the optimal value translates into a variation of species abundances related to trait values, the mean value of trait in communities (Community-Weighted Mean, CWM) and their variance (CWV) (Garnier et al. 2016) are expected to reflect local optimal trait value and filtering intensity, respectively (Cingolani et al. 2007, Violle et al. 2007, Enquist et al. 2015, Borgy et al. 2017a). However, a clear relationship between trait-based statistics and the parameters of environmental filtering (“CWM-optimality” hypothesis, Muscarella and Uriarte 2016) may not always hold.

**Figure 1.**
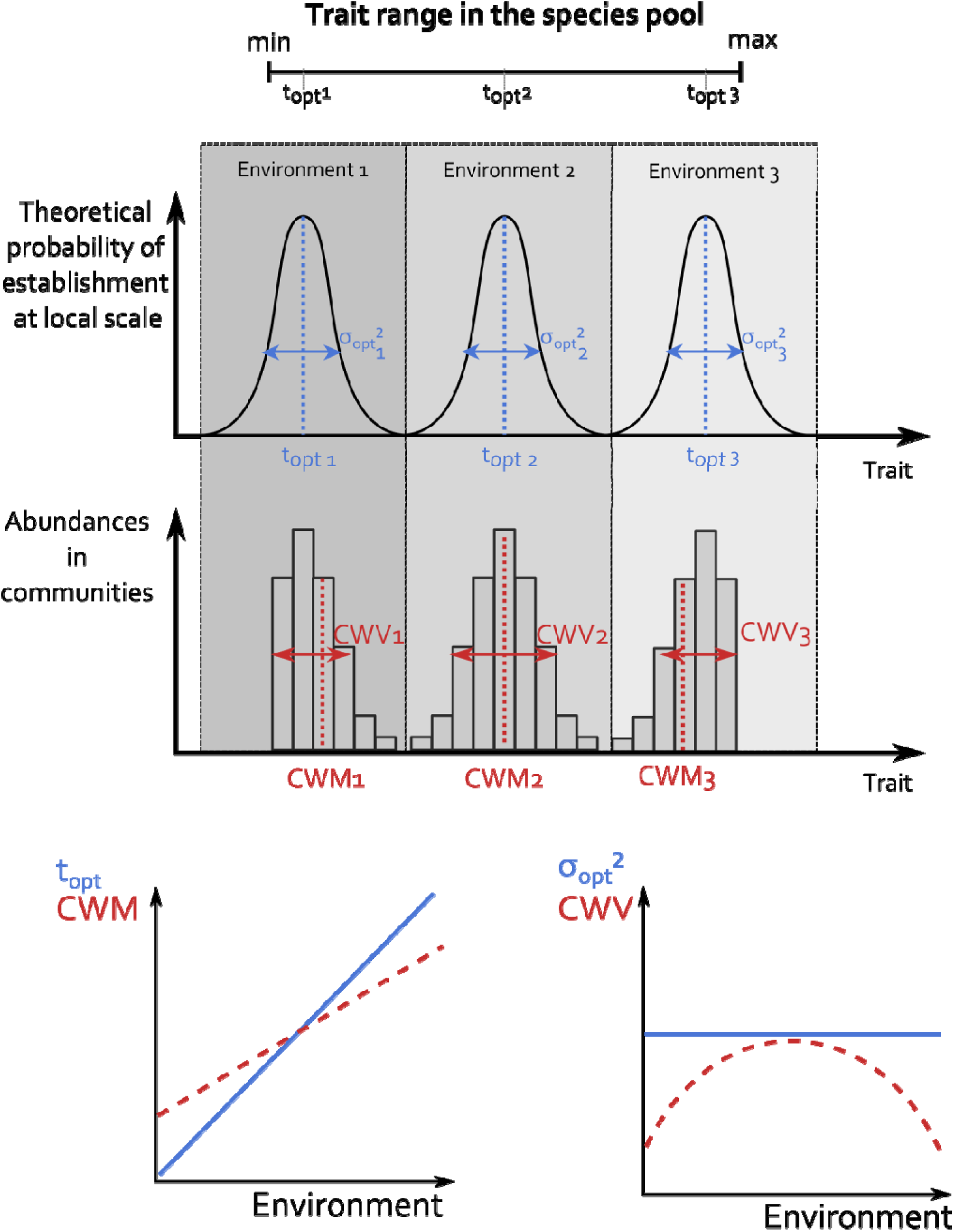
Departure of CWM and CWV from the parameters of environmental filtering, *t*_*opt*_ and *σ*_*opt*_^*2*^, respectively, due to trait limits in the species pool. The trait distribution in communities (histograms) reflects the joint influence of trait range limits among immigrants from the species pool (top horizontal black line), and of a Gaussian environmental filter determining immigrant establishment success with mean *t*_*opt*_ (dashed blue lines) and standard deviation *σ*_*opt*_^*2*^ (blue horizontal arrows) in specific environments (grey rectangles). The dashed red lines represent the observed Community Weighted Mean (CWM) values in each community. CWM deviates from *t*_*opt*_ when closer to the limits of the trait range in the species pool because of the bounded species pool’s trait range. The range of observed CWM values (red segment) is then smaller than the one of *t*_*opt*_ values as shown in the CWM ~ Environment plot. Similarly, while *σ*_*opt*_^*2*^, which represents the environmental filtering intensity, remains constant over the environment gradient, CWV, depicted by the horizontal red arrows, decreases when approaching environment selecting for trait values closed to the species pool boundaries. The hump-shaped relationship between realized CWV and the environment thus represents the influence of the trait range limits and not a more intense filtering at the extremes of the gradient.

In extreme environments, more intense environmental filtering due to local constraints is commonly hypothesized (Weiher et al. 1998, Callaway et al. 2002, Cornwell et al. 2006), but the filtered trait values can also be closer to regional trait range limits. A reduction of variance in extreme environments can thus be allotted to either local environmental filtering or to larger-scale and longer-term constraints leading to a restricted trait variation among immigrants. Regional trait range limits should yield a decrease in local trait variance at the extremes of an environmental gradient and therefore entail a hump-shaped variation of CWV across the environmental gradient, even when the intensity of environmental filtering is constant throughout the gradient (Fig. 1). TGBE can also originate phenomenological relationships between CWM and CWV because of the local convergence induced by the species pool limited trait range. Such hump-shaped patterns between CWM and CWV have been reported previously (Dias et al. 2013), and can reflect the influence of TGBE in real data. A major issue is then to determine whether lower trait variance in extreme environments reflects more intense filtering or the influence of trait limits at a regional scale. To solve the issue, we propose an inference approach that explicitly estimates the influences of regional trait range limits and local environmental filtering.

We investigated TGBEs in the context of a spatially-implicit model of community assembly representing how immigration from a species pool and local environmental filtering jointly shape local community composition (*ecolottery* package, Munoz et al. 2018) (Fig. 1). Environmental filtering is modeled as a Gaussian function determining the successful establishment of immigrants and thus defines a decrease of the performance of species around an optimum trait value, the intensity of the filtering being the standard deviation of the function (Webb et al. 2010, Shipley 2010, Enquist et al. 2015). An environmental gradient can then be viewed as a gradient of distinct optima imposed by distinct local environmental filters. When trait range limits among immigrants constrain the functional range in community composition, we expect reduced variance and a skewed local distribution with CWM deviating from optimal trait value (Fig. 1). We used the model to simulate community composition with explicit environmental filtering along an environmental gradient, with and without variation of filtering intensity. It illustrates how TGBEs can arise. In addition, we propose an Approximate Bayesian Computation approach based on intensive simulations of community composition to get an unbiased estimate of the optimum and intensity of environmental filtering, while controlling for the influence of TGBE. This powerful and mechanistic approach allows comparing the outputs of our community assembly model, with different sets of parameters related to distinct processes, to the local trait patterns observed in a given community dataset, so as to unravel the causes originating them (Csilléry et al. 2010, Munoz et al. 2018). We applied the approach to examine TGBE and environmental filtering in alpine plant communities along a gradient of snow cover duration in the French Alps (Choler, 2005).

## Material and Methods

### Framework of community assembly

Immigrants drawn from a species pool establish and persist in a community depending on environmental filtering (Fig. 1). Each individual displays a synthetic fitness-related trait value, *t*, and the probability of successful immigration decreases as *t* departs from an optimal trait value *t*_*opt*_ depending on local environmental conditions (Shipley 2010). We used a Gaussian function of *t* with mean *t*_*opt*_ and standard deviation *σ*_*opt*_ to represent this filtering. *σ*_*opt*_ depicts the intensity of environmental filtering: the smaller *σ*_*opt*_, the narrower the extent of trait values allowing immigration in the local community (Munoz et al. 2018). Each community is then assigned *t*_*opt*_ and *σ*_*opt*_ values characterizing local environmental filtering.

Our main objective is to disentangle the influence of (i) trait range limits in the species pool, denoted as *a* for the lower and *b* for the upper limit, and (ii) the parameters of environmental filtering denoted as *t*_*opt*_ and *σ*_*opt*_, on the distribution of trait values in local communities. When *t*_*opt*_ is close to *a*, we expected that the distribution of trait values in the local community is limited below *a* (Fig. 1), and conversely when *t*_*opt*_ is close to *b*. In the following, we present the consequences of the regional trait limits on (i) the calculation of the first four moments (Enquist et al. 2015) of the local trait distribution, and (ii) how these moments vary across communities along an environmental gradient.

### Community-level trait based statistics

Synthetic trait-based statistics are commonly used to characterize the functional response of communities. The two first moments of the distribution of trait values in a community, namely, the community weighted mean (CWM) and community weighted variance (CWV), are commonly used to analyze the functional structure of communities while the two following moments, community weighted skewness (CWS) and community weighted kurtosis (CWK) are more rarely considered (Enquist et al. 2015, Gross et al. 2017). The first four moments are expected to be influenced, among other processes, by environmental filtering and are often used for the inference of filtering (Shipley 2010, Enquist et al. 2015, Loranger et al. 2018). With a Gaussian environmental filtering (Fig. 1), we expect CWM and CWV to equal *t*_*opt*_ and *σ*_*opt*_, respectively. As a measure of ‘peakedness’, CWK should also increase with decreasing *σ*_*opt*_ (Enquist et al. 2015, Gross et al. 2017). If the environmental filter is symmetrical, as considered here (Fig. 1), local CWS is not expected to deviate from 0.

When the trait range in the species pool is bounded and when the environment selects for trait values close to these boundaries, the local distribution of trait values is bounded beyond the limits of the pool, and is asymmetrical (Fig. 1). This asymmetry should entail a shift in CWM to larger values if the closer trait limit in the species pool is the lower boundary and to lower values if the closer limit is the upper boundary (Fig. 1). In addition, the trait limits should further reduce the range of values in local communities and thus reduce CWV (Fig. 1), increase CWK and increase CWS in absolute value when *t*_*opt*_ is closer to the limits. In Appendix S1, we provide the mathematical formulas of the four moments, as a function of *t*_*opt*_, *σ*_*opt*_, and of trait range limits *a* and *b*, in a simple case where regional trait abundances are uniformly distributed between *a* and *b*.

### Simulation of communities with environmental filtering and trait range limits

We used a coalescent-based algorithm (package *ecolottery* in R language, Munoz et al. 2018) to simulate community assembly with migrants drawn from a species pool and subject to a Gaussian environmental filtering. The coalescent-based approach reconstructs the shared ancestry of coexisting individuals (i.e., their genealogy) at present without simulating complete community dynamics from an initial state through time. The topology of the genealogy depends on immigration, environmental filtering, and demographic stochasticity (Munoz et al. 2018). We considered two types of species pools with either a uniform or a log-series distribution of abundances. Results were comparable with both distributions, and subsequent analyses will concern the case of uniform abundances only. A uniform pool includes 100 species with 1,000 individuals per species, hence a total of 100,000 candidate immigrants. Species trait values *t*_*i*_ were drawn from a uniform distribution between either *a* = 0 and *b* = 1 (trait range = 1), or *a* = 0 and *b* = 2 (trait range = 2). We varied the range of trait values to assess the relative influence of filtering intensity and trait range. We also simulated a set of communities with intraspecific variation, i.e., with a standard deviation of trait values per species set to *σ*_*i*_ = 0.1 in the species pool. The environmental filtering function determined the probability *p* of establishment of an individual with a trait value *t* according the following function: 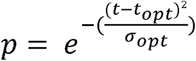 (Fig. 1). We set the intensity of environmental filtering, ruled by the parameter *σ*_*opt*_, to either 0.25 or 0.05, to represent weak and intense environmental filtering, respectively, compared to the regional range of trait values varying between 0 and 1. For a given species pool, we simulated *n* = 100 communities, each including *J* = 500 individuals, with varying *t*_*opt*_ values randomly drawn between *a* and *b*. The variation of *t*_*opt*_ represents a variation of optimal values along the environmental gradient.

We also considered another set of simulations where *σ*_*opt*_ varied along the gradient, with minimum values of *σ*_*opt*_ = 0.05 at the extremes *a* and *b* towards a maximum of *σ*_*opt*_ = 0.25 in the middle of the gradient. In this case, environmental filtering was more intense at the extremes of the gradient. We therefore designed two sets of simulated communities undergoing a fixed and varying environmental filtering, respectively. From these simulated data, local weighted moments were calculated and the environmental filtering parameters 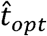 and 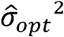 were estimated (see below). A repeatable example of community simulation is provided in Appendix S2.

### ABC estimation of parameters of environmental filtering

We performed an Approximate Bayesian Computation (ABC) analysis (Csilléry et al. 2010, *coalesc_abc* function in *ecolottery* R package) to estimate the parameters 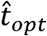 and 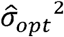 of environmental filtering from a given community composition. ABC provides posterior distributions of parameter estimates by comparing some summary statistics in communities simulated over a broad range of *t*_*opt*_ and *σ*_*opt*_ values, to the same summary statistic values in the given community (Csilléry et al. 2010). In our case, the summary statistics were metrics of taxonomic (richness and Shannon diversity) and functional (CWM, CWV, CWS and CWK) composition of a community. Many communities were simulated in ABC analysis using the same coalescent-based algorithm presented above (package *ecolottery* in R language, Munoz et al. 2018). In any case, simulated communities received immigrants from the same species pool. We also considered an alternative analysis where the summary statistics included functional dispersion (Laliberté and Legendre 2010) and Rao’s quadratic entropy (Botta-Dukát 2005) instead of CWV, CWS and CWK (Appendix S10). Insofar as species pool composition was known, its trait range limits *a* and *b* were fixed based on the upper and lower trait range limits in the complete species pool. However, we also devised a case where the trait range limits and the species pool composition were based on the sum of observed communities (Appendix S11). The median values of 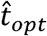 and 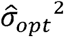 in posterior distributions were compared to observed CWM and CWV values, respectively.

We performed ABC analysis on each of the simulated community presented above, to get a cross-validation of estimated 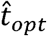 and 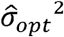 values for simulated data with known *t*_*opt*_ and *σ*_*opt*_^*2*^ values. We also compared CWM and CWV in communities to *t*_*opt*_ and *σ*_*opt*_^*2*^. Figures 2b and 2d represent the variation in ABC estimates along a gradient of *t*_*opt*_ values. For simulations with fixed *σ*_*opt*_^*2*^, any variation in CWV at the extremes was expected to reveal an influence of regional trait limits only (TGBE). Conversely, we expected decreasing 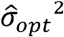 at the extremes of the gradient of *t*_*opt*_, for the set of simulations where *σ*_*opt*_^*2*^ was indeed smaller at the extremes. The 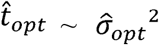 relationship was also compared to the *CWM* ~ *CWV* relationship, to check the consistency of the variation in estimated environmental filtering parameters with phenomenological patterns of functional convergence measured with CWV (Appendix S3).

**Figure 2.**
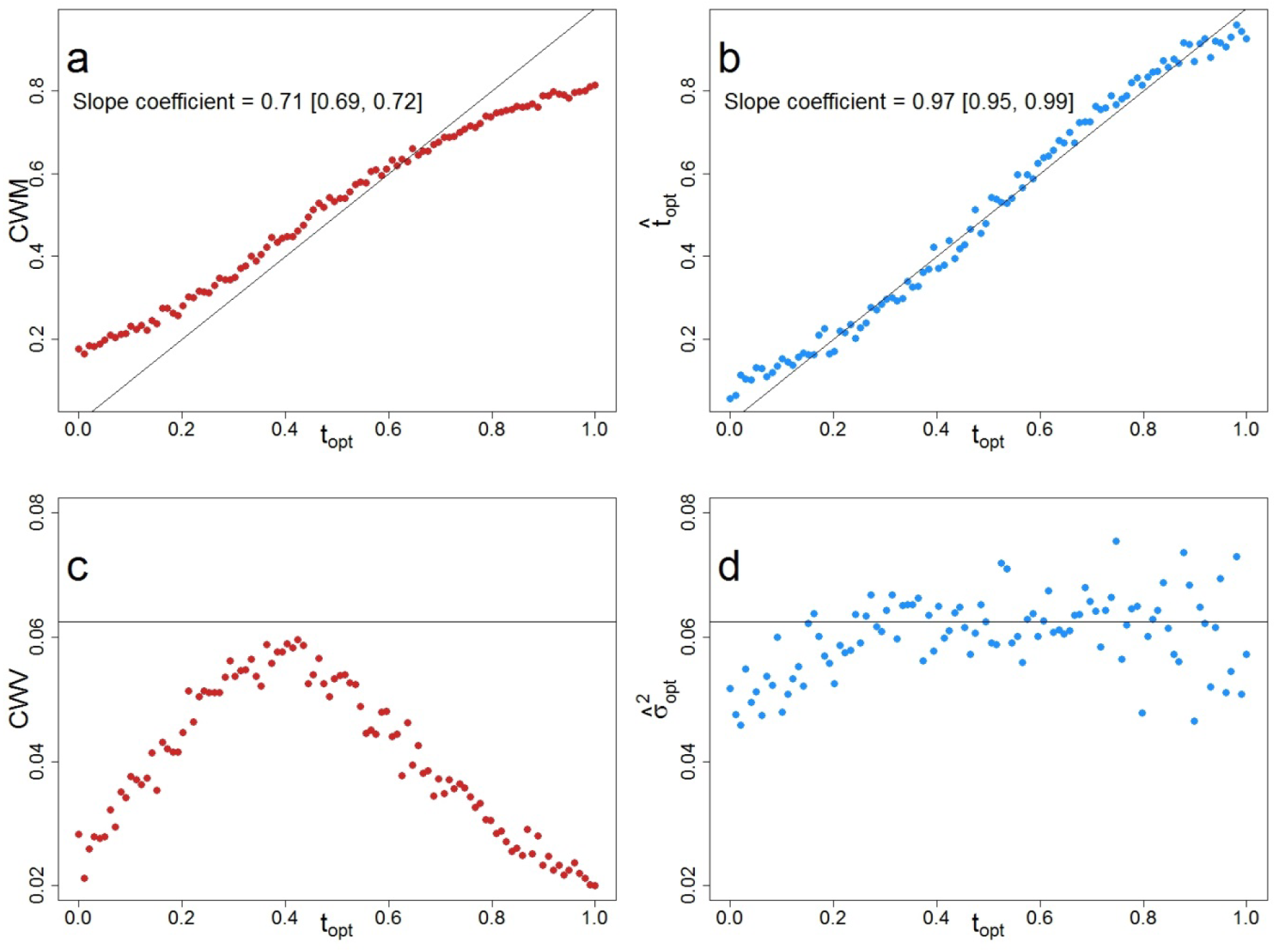
Variation in CWM and CWV values (left, red color), and of estimated 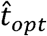 and 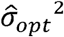 (right, blue color), for simulated communities along *t*_*opt*_ gradient. Communities were simulated with constant environmental filtering (*σ*_*opt*_ = 0.25), uniform distribution of trait values and uniform abundances in the species pool. Top figures (a) and (b) represent CWM and 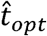, and figures (c) and (d) represent CWV and 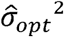. The 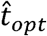 and 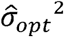 values were obtained with the ABC approach and correctly estimated the *t*_*opt*_ and *σ*_*opt*_^*2*^ values (b and d). Conversely, CWM departed from *t*_*opt*_ and CWV was below *σ*_*opt*_^*2*^ when the influence of trait range limits increased at the extremes. The black solid line represents equality of CWM and CWV to the parameters of environmental filtering (*t*_*opt*_ and *σ*_*opt*_^*2*^, respectively). Slope coefficients and the associated confidence intervals of the linear regression equations between CWM and *t*_*opt*_ are displayed in panel (a) and (b). The mean of the difference between *σ*_*opt*_^*2*^ and CWV (c) is twice higher than for the difference between *σ*_*opt*_^*2*^ and 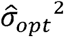 (d) (respectively 2.23e-2 and 8.91e-3).

### Application to alpine plant communities

We analyzed the variation in functional composition of plant communities along a gradient of snow cover duration in alpine grassland vegetation (Choler 2005). This gradient ranged from 140 to 210 days of snow cover in 1998. The alpine vegetation dataset (*aravo* in ade4 R package) includes 75 communities for a total of 82 species, located between 2700 and 2750 meters in French Alps. This vegetation undergoes harsh high-elevation conditions but also covers a broad environmental gradient of duration of snow cover, due to topographical and microclimatic heterogeneity (Choler 2005). The gradient determines varying abiotic stress and length of growing season, and thus largely influences functional trait variation among communities, such as leaf nitrogen concentration on a mass basis (N_mass_) and specific leaf area (SLA) (Choler 2005), which are two foliar traits characterizing the resource acquisition-conservation tradeoff in plants (Garnier et al. 2016). Long snow cover protects from freezing stress but reduces the length of growing season, which should favor resource-acquisitive plants, relatively to the local species pool, with higher N_mass_ and SLA. On the contrary, short snow cover increases exposure to wind and frost while increasing length of growing season, which should, in this specific context, favor resource-conservative plants with lower N_mass_ and SLA (Choler 2005).

We estimated parameters of environmental filtering *t*_*opt*_ and *σ*_*opt*_ for foliar traits in this dataset, and examined their variation along the gradient of snow cover duration. The species pool used in ABC analysis was built from the species present in all the observed communities.

## Results

### TGBE in simulated communities

We simulated communities along an environmental gradient with different *t*_*opt*_ values but constant filtering intensity *σ*_*opt*_^*2*^ (Fig. 2). The variations in CWM and CWV illustrate the influence of TGBE. First, CWM went below *t*_*opt*_ when closer to the upper limit of trait range, and above *t*_*opt*_ when closer to the lower limit (Fig. 2a). The observed range of CWM values was thereby smaller than the range of *t*_*opt*_. Second, we found a hump-shaped variation in CWV, with lower values at the extremes of the *t*_*opt*_ gradient (Fig. 2c). CWS and CWK also varied along the *t*_*opt*_ gradient with a decrease in CWS and an increase in CWK towards the extremes (Appendix S5). Because filtering intensity was set constant, the reduction of CWV at the extremes, and the respective variations of CWS and CWK, was attributable to the influence of trait range limits in the species pool (Fig. 1). We obtained consistent results under more intense but constant environmental filtering (σ_opt_ = 0.05, Appendix S4, more contrasted), with intraspecific variability (Appendix S6, σ = 0.1), with log-series distribution of regional abundances (Appendix S7) and when using the sum of observed communities as a species pool (Appendix S11).

We expected the influence of TGBE to extend farther from the extremes when *σ*_*opt*_^*2*^ was larger for a fixed range [*a*; *b*]. The extent of the influence of regional trait limits was thereby expected to depend on the intensity of local filtering relatively to trait range [*a*; *b*]. Appendix S8 shows how the ratio of *σ*_*opt*_^*2*^ and trait range influences the deviation of CWM from *t*_opt_. It shows that the ratio of trait range (*b* - *a*) and filtering intensity (*σ*_*opt*_^*2*^) determines the influence of TGBE along the gradient. For instance, *σ*_*opt*_ = 0.5 and [0; 1] trait range gives the same deviation than σ_*opt*_ = 1 and [0; 2] trait range.

### Deciphering environmental filtering and TGBE in extreme environments

In communities where filtering intensity was set constant, we obtained unbiased estimation of *t*_*opt*_ (Fig. 2b, slope coefficient of the regression between 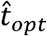 and *t*_*opt*_ = 0.97), and unbiased and constant estimation of *σ*_*opt*_, while there was variation in CWV due to TGBE (Fig. 2d). Indeed, the square distance between *σ*_*opt*_^*2*^ and 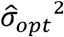 was, in average, twice low over the *t*_*opt*_ gradient (Fig. 2d) than the square distance between *σ*_*opt*_^*2*^ and CWV (Fig. 2c) (8.91e-3 and 2.23e-2 respectively). When using other metrics than CWV to evaluate local functional convergence and to estimate *t*_*opt*_ and *σ*_*opt*_^*2*^, namely functional dispersion and Rao’s quadratic entropy, we obtained similar results with significant quadratic relationships between these metrics and *t*_*opt*_ along *t*_*opt*_ gradient while the environmental filtering intensity remained constant (Appendix S10). In addition, we simulated an environmental gradient where filtering was more intense at the extremes (i.e., smaller *σ*_*opt*_ value, black line on Fig. 3a and 3b). Figure 3d shows that the estimated value of *σ*_*opt*_ followed the expected variation of filtering intensity. In this case, CWV also displayed a hump-shaped pattern along the gradient, similar to Figure 2c, but here this was due to both regional trait limits and actual variation in filtering intensity.

**Figure 3.**
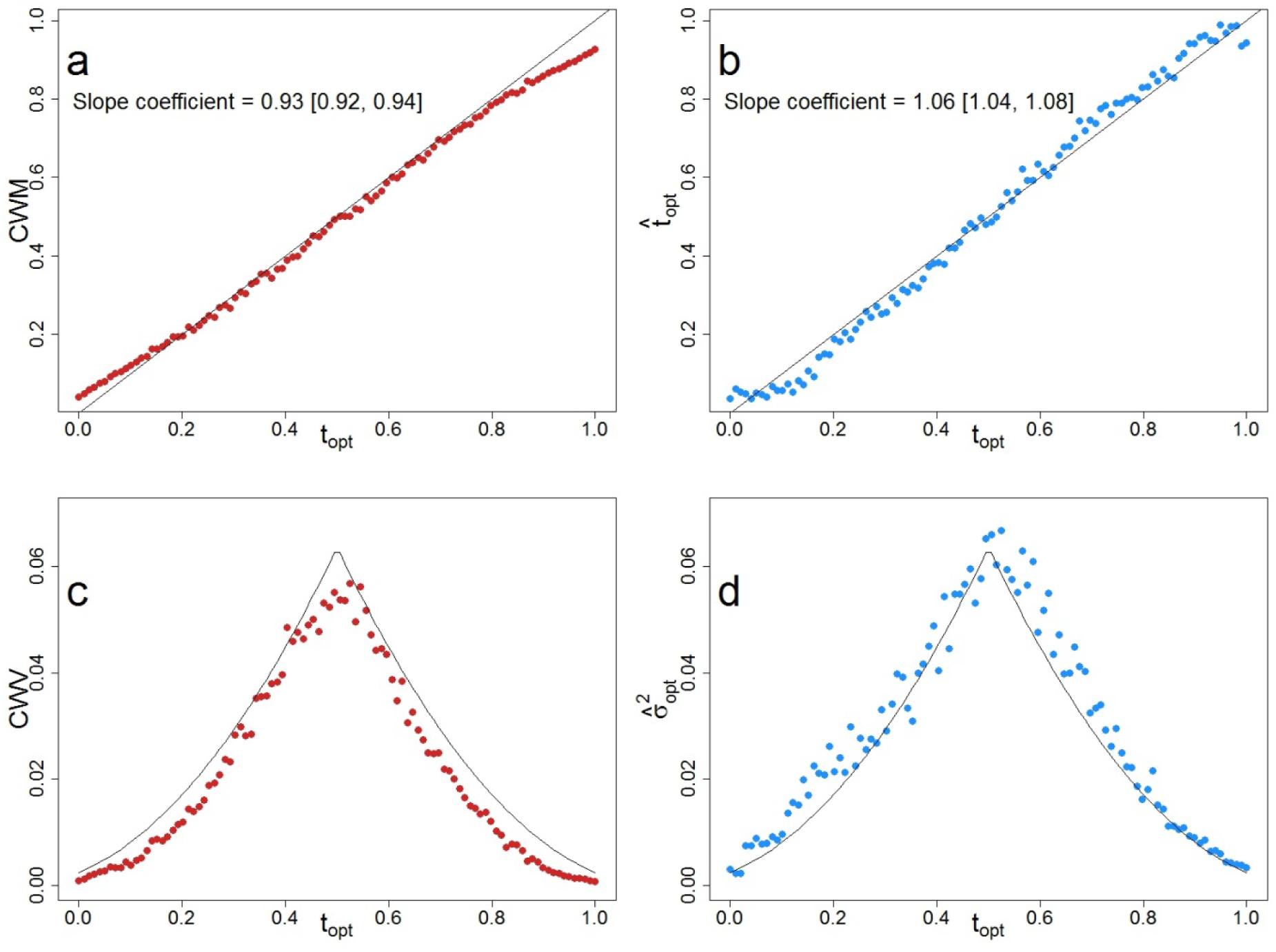
Variation in CWM and CWV (left, red color), and in estimated 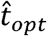 and 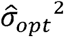 (right, blue color) along the *t*_*opt*_ gradient, with increasing intensity of environmental filtering at the extremes of the gradient. Top figures (a) and (b) represent CWM and 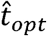, and figures (c) and (d) represent CWV and 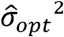. The estimation of parameters 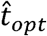 and 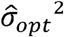, obtained with the ABC approach, acknowledges the effect of trait range limits, and departs from CWM and CWV, respectively when the influence of the trait range limits increases at the extremes. The black solid line represents equality of CWM and CWV to the parameters of environmental filtering (*t*_*opt*_ and σ_opt_, respectively). Slope coefficients and the associated confidence intervals of the linear regression equations between CWM and *t*_*opt*_ are displayed in panel (a) and (b). The mean of the difference between *σ*_*opt*_^*2*^ and CWV (c) and between *σ*_*opt*_^*2*^ and 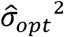 (d) is comparable but lower for the latter case (respectively 4.08 e-2 and 3.37e-2).

Therefore, the variation in CWV could not inform on the respective influences of environmental filtering and trait range limits in the pool (Fig. 1c, Fig. 3c), while the ABC-based estimation of *σ*_*opt*_^*2*^ allowed grasping the specific influence of environmental filtering.

### TGBE and environmental filtering in alpine plant communities

We estimated *t*_*opt*_ and *σ*_*opt*_^*2*^, and the variations in CWV and CWM values of foliar traits in alpine plant communities (Figs. 4 & 5). As expected with TGBE, CWM departed from estimated 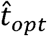 in extreme environmental conditions, and the range of *t*_*opt*_ values was larger than the range of CWM values (Fig. 4ab & 5ab). CWV decreased at lowest duration (great exposure to cold) for both SLA and N_mass_ and at highest duration (short vegetative period) of snow cover for N_mass_ only (Fig. 4). On the contrary, ABC-based estimations showed that 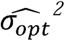 did not vary along the snow cover gradient (Fig. 4cd & 5cd). Except for SLA at long snow cover duration (Fig. 5cd), 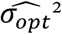 was larger than the corresponding CWV.

**Figure 4.**
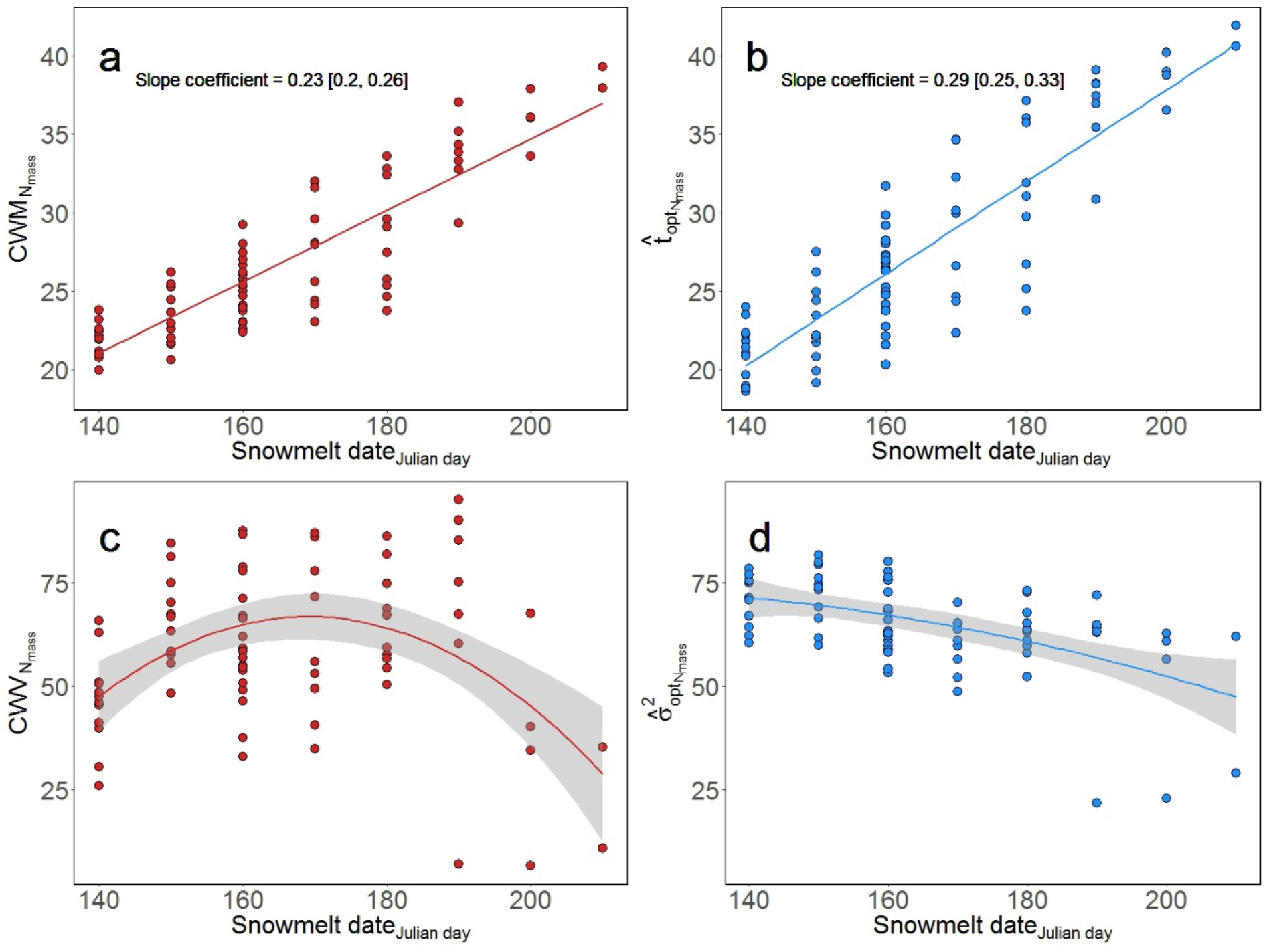
Relationships of CWV (red color) and estimated *σ*_*opt*_^*2*^ (blue color) for the leaf nitrogen content on a mass basis (N_mass_, panel a) and the specific leaf area (SLA, panel b), according to the gradient of snow cover melting date (in Julian days, abscissa). Linear regressions were fitted for each variable against the snowmelt date in panels a and c. While both highly significant, the slope term was higher for with the estimated 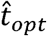 (slope = 0.29) than with the CWM (slope = 0.23). For the panels c and d, a quadratic regression between CWV and snowmelt date was significant while the quadratic term became non-significant with a 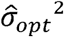 is measured in mg[N]/mg.

**Figure 5.**
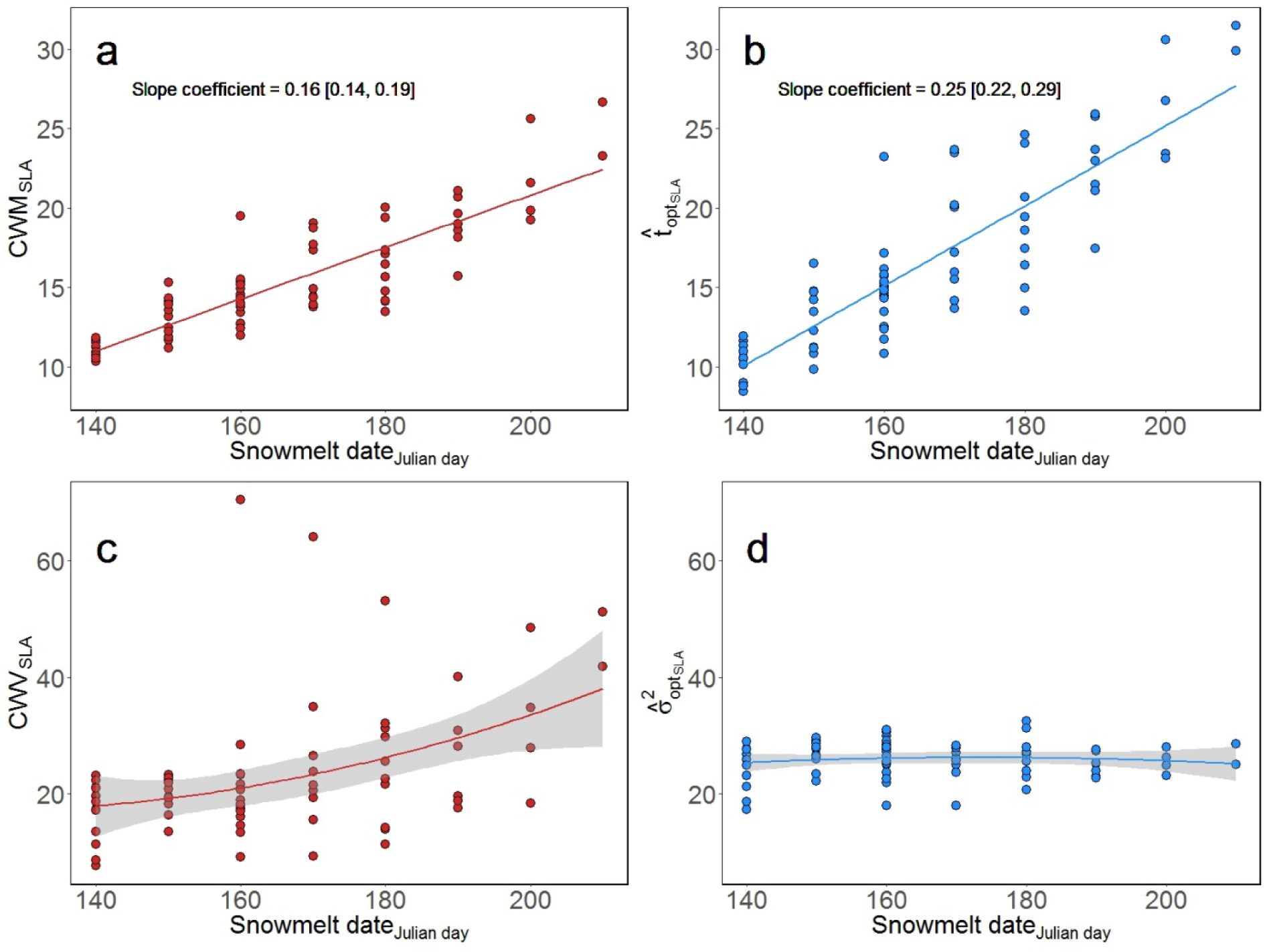
Relationships of CWV (red color) and estimated *σ*_*opt*_^*2*^ (blue color) for the specific leaf area (SLA), according to the gradient of snow cover melting date (in Julian days, abscissa). Linear regressions were fitted for each variable against the snowmelt date in panels a and c. While both highly significant, the slope term was higher for with the estimated 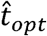 (slope = 0.25) than with the CWM (slope = 0.16). For both panels c and d, both quadratic regressions between CWV and a 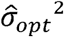 with snowmelt date were non-significant. SLA is measured in m^2^/kg.

In addition, departure of community weighted skewness (CWS) from 0 reflected the influence of regional trait limits and asymmetry in local trait distribution, as observed in simulated communities with constant *σ*_*opt*_ (Appendix S5). In alpine plant communities, increasingly negative community weighted skewness (CWS) with increasing snow cover duration in alpine vegetation was consistent with an influence of an upper trait limit on the local distribution of N_mass_ and SLA at longest snow cover duration (Appendix S9).

## Discussion

In ecology and biogeography, trait-gradient analyses examine the functional trait distributions in communities to characterize community responses along environmental gradients (Ackerly and Cornwell 2007, Lepš et al. 2011, Garnier et al. 2016, Borgy et al. 2017a). Here we showed that a reduced variance of the local trait distribution, i.e., trait convergence, can reflect a combined influence of local environmental constraints within the community and of a bounded trait distribution in the regional species pool. These two influences need to be disentangled in order to identify the specific role of local environmental filtering. However, while much emphasis has been put on the idea that environmental filtering can be more intense at the extremes of environmental gradients (Weiher et al. 2011), far less attention has been devoted to how the functional composition of species pools influences local community composition (Spasojevic et al. 2018). To address the issue, we used a simulation-based, Approximate Bayesian Computation (ABC) approach (*ecolottery* package, Munoz et al. 2018). By explicitly modelling immigration and environmental filtering, the approach allows separating out the influence of constraints on trait distributions at species pool and local community levels. With this approach, we can obtain unbiased estimation of *t*_*opt*_ and *σ*_*opt*_ in simulated communities along gradients. The mid-domain effect is a better-known example of the influence of regional limits (of species niches and distributions) influencing local taxonomic diversity at the extremes of gradients (in geographical, Colwell and Lees 2000, or environmental space, Letten et al. 2013). The TGBE issue presented here extends this perspective to examine how trait range limits in species pools influence functional composition in local communities. We discuss the consequences of TGBE for trait-based approaches in functional ecology, community ecology and (functional) biogeography.

Environmental filtering is often viewed as a humped filtering function along a niche axis, similar to a Gaussian function with optimal value *t*_*opt*_ and filtering intensity *σ*_*opt*_. Although environmental filtering generally concerns the influence of abiotic constraints (Kraft et al. 2015), the framework proposed here can apply to any filtering around an optimal trait value *t*_*opt*_ conferring, e.g., greater competitive ability (Mayfield and Levine 2010), better colonization or chances of establishment (Ehrlén and Eriksson 2000, Bernard-Verdier et al. 2012). The current paradigm in functional ecology is that community weighed mean (CWM) is a proxy for *t*_*opt*_, the “CWM-optimality” hypothesis (Muscarella and Uriarte 2016), and that community weighed variance (CWV) is a proxy for *σ*_*opt*_^*2*^ under environmental filtering. The “CWM-optimality” hypothesis found some support in recent studies linking the distance between species’ traits and CWM to species’ abundances for single traits (Umaňa et al. 2015) or multivariate measures (Muscarella and Uriarte 2016), but was challenged in other contexts (Mitchell et al. 2017, Laughlin et al. 2018). CWM can be disconnected from *t*_*opt*_ when stabilizing mechanisms such as competitive interactions and limiting similarity break the linkage of trait values with fitness differences (Chesson 2000, Adler et al. 2013), or when neutral stochastic dynamics affect species abundance independently from trait values (Hubbell 2001). Here we challenge the CWM-optimality hypothesis by demonstrating that CWM and CWV can depart from *t*_*opt*_ and *σ*_*opt*_^*2*^, respectively, when the local distribution is bounded due to trait range limits in the pool of immigrants. The distribution of trait values in the regional species pool therefore influences local community assembly (Patrick and Brown 2018, Spasojevic et al. 2018) and can challenge the CWM-optimality hypothesis by preventing CWM to reach the optimum for certain environments. It is likely that trait range limits of the species pool are reached in extreme environments, i.e. trait values required for persistence are not possible, due to physiological limits or evolutionary history (Koch et al. 2004, Alpert 2005). It is essential to distinguish the respective signatures of local environmental filtering and of processes driving the functional composition of species pools at a larger scale and over a long term (Jiménez-Alfaro et al. 2018). Consequently, identifying TGBEs means determining the specific influence of local community assembly amidst the influence of large-scale and long-term evolutionary legacy (Lessard et al. 2016).

We found that TGBE can be responsible of a hump-shaped variation in CWV along environmental gradients even when the intensity of environmental filtering is constant (Fig. 2c). TGBE also generated a hump-shaped relationship between CWV and CWM (Appendix S3), similar to patterns reported in a previous study (Dias et al. 2013). Although a link between CWM and CWV (or similar functional diversity metrics) can represent a statistical artifact (Ricotta and Moretti 2011, Dias et al. 2013), our study also shows that TGBE can yield this relationship. The analysis of alpine plant communities illustrated trait variance reduction in extreme environmental conditions (Fig. 4 & 5), while the estimated 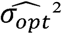 did not show reduction. Variance reduction could thus be due to TGBE and not to more intense environmental filtering in these alpine plant communities (Fig. 4). Similarly, the 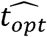-environment relationships had a steeper slope than the CWM-environment relationship (Fig. 2, Fig. 4ab & Fig. 5ab), suggesting that CWM did not represent optimal trait values all along the environmental gradient.

We have proposed a spatially-implicit framework of community assembly acknowledging immigration from a species pool and local environmental filtering (Munoz et al. 2018). The definition of the pool is flexible and several options have been proposed, either based on a regional list of species (Zobel 1997), on the complete composition of a metacommunity (Leibold et al. 2004), or on a spatially restricted source of dispersers (Lessard et al. 2016). The pool can represent an external forcing based on long-term and large-scale regional dynamics (top-down perspective as in Hubbell 2001) or reflect the emergent composition of available immigrants in a metacommunity (bottom-up perspective, Leibold et al. 2004, Mittelbach and Schemske 2015). In both cases, its composition illustrates the influence of long-term assembly dynamics across communities in a specific area, and its boundaries represent the limits imposed by these processes. In the present analyses, while we simulated and used the composition of complete species pools in ABC analyses of simulated communities, the species pool of alpine communities was based on the sum of sampled communities (see in Appendix S11 the results for simulations with a species pool based on the sum of sampled communities). The composition and the relative abundances considered in the reference species pool can greatly influence analyses of community assembly dynamics (Lessard et al. 2011). Dark diversity, representing the species that are absent from the pool but could contribute to immigration and community assembly (Pärtel et al. 2011), can extend trait range limits in the reference species pool. Further investigation of the influence of trait range limits with different definitions of the species pool should help address under which conditions TGBE can be reliably detected. Furthermore, the influence of the shape of the trait distribution in the pool should be addressed in more details in the future (Spasojevic et al. 2018) and appears essential since it can vary from a biogeographical context to another even though local environmental filtering can operate in a similar way. For sake of simplicity, we considered a uniform distribution of trait values among species at regional scale, and two types of distribution of regional abundances, uniform and log-series. Even though the results were robust to some variation in these parameters, further investigation of the sensitivity of the model will be needed. Lastly, we defined environmental filtering in our study as a Gaussian function around a single optimum (Shipley 2010). However, other filtering functions, such as disruptive filtering with two modes yielding trait divergence (Loranger et al. 2018), could be considered to study trait patterns at the community level, and are already implemented in *ecolottery* R package (Munoz et al. 2018).

Independently from the assumptions mentioned above, the way CWM and CWV deviate from *t*_*opt*_ and *σ*_*opt*_^*2*^ due to TGBE depends on the ratio between the trait range limits and the strength of local environmental filtering along a gradient (Appendix S8). In a biogeographical perspective, a physiological trait ~ environment relationship could yield different patterns of CWM and CWV variation across regions where distinct biogeographical histories entailed different range limits (Forrestel et al. 2017). Moreover, for a given regional species pool, the influence of TGBE should change depending on the strength of local environmental filtering. Therefore, when the filtering acting on a specific trait is strong, the deviation should concern only communities closest to the extremes. The influence of TGBE on trait ~ environment relationship can also differ across functional traits, depending on the nature of underlying filters acting on different traits (Borgy et al. 2017b). The detection and influence of TGBE will therefore be dependent upon the interplay of biogeographical history and the local mechanisms filtering, with certain intensity, trait values.

ABC-based estimation of environmental filtering relies on simulating and comparing basic statistics that summarize the observed and simulated trait distributions. The moments of local trait distributions can be used as summary statistics to infer the trait-based assembly processes, as advocated by the Trait Driver Theory (TDT) (Enquist et al. 2015). While much emphasis has been put on analyzing the two first moments CWM and CWV, TDT underlines that the next moments, skewness (CWS) and kurtosis (CWK), also convey insights on assembly dynamics. Gross et al. (2017) emphasized that CWS and CWK allow better characterizing the coexistence of multiple functional strategies beyond the influence of a single optimum. We showed that TGBE strongly impacts CWV variations (Figs. 1, 2c and 2d) but also other moments (Appendices S5 and S9). As a consequence, applying TDT along gradients also probably implies addressing TGBE issues. Community-level metrics are more and more used to characterize the functional composition of communities of plants (Violle et al. 2007), but also other organisms (e.g., Newbold et al. 2012, Fierer et al. 2014, Pey et al. 2014). We stress here that these metrics should not be viewed as direct proxies of underlying assembly processes, especially in harsh environmental conditions that are the focus of much research and where TGBE more likely occurs. Furthermore, acknowledging intraspecific variation in trait-based community analyses has gained much momentum in recent years (Lepš et al. 2011, Violle et al. 2012, Siefert et al. 2015). Having intraspecific trait variation could extend beyond the trait limits of a pool defined based on trait values averaged at species level (Violle et al. 2012), which should affect associated trait range limits and therefore TGBE. Our individual-based modelling framework can acknowledge the influence of intraspecific trait variation in community dynamics (Appendix S6), but these data are mostly unavailable at large spatial scales of functional biogeography, so that trait values averaged at species level are still mainly used in practice (Borgy et al. 2017b).

Community-level trait metrics are common currencies for functional biogeography (Violle et al. 2014). They can be used to elucidate the drivers of taxonomic diversity patterns (Lamanna et al. 2014) as well as to target conservation areas (Violle et al. 2017) or to map and predict ecosystem properties from landscape to regional and global scales (Violle et al. 2015). The approach is primarily based on the “CWM-optimality” hypothesis (Muscarella and Uriarte 2016), and the idea that CWV reflects the intensity of the local environmental filtering. Other processes can affect local community assembly and functional composition (Hubbell 2001, Levine and Murrell 2003, Mayfield and Levine 2010, Muscarella and Uriarte 2016), and our work further underlines that the functional composition of the species pool providing immigrants is influential. Taking into account the functional diversity of the species pool, and acknowledging the underlying biogeographical and evolutionary dynamics, is an important issue that has only recently come to focus (Patrick and Brown 2018, Spasojevic et al. 2018). TGBE shows the need to better integrate local and regional dynamics when examining the functional composition of local communities. Therefore, ecologists need to be aware of TGBE when interpreting patterns of functional composition and their causes, notably at the extremes of environmental gradients.

## Declarations

## Acknowledgements

This study was supported by the European Research Council (ERC) Starting Grant Project ‘Ecophysiological and biophysical constraints on domestication in crop plants’ (Grant ERC-StG-2014-639706-CONSTRAINTS and by the French Foundation for Research on Biodiversity (FRB; www.fondationbiodiversite.fr) in the context of the CESAB project ‘Causes and consequences of functional rarity from local to global scales’ (FREE).

## Data Accessibility

R code to generate the simulated data is provided in Appendix S2 and is archived on a Zenodo URL link: https://zenodo.org/record/1165400.

The *aravo* dataset describing alpine plant communities is available in the ade4 R package and is described in Choler (2005).

## Supporting information

## Appendix S1.

Moments of the located trait distribution.

With a uniform distribution of species trait values in the species pool ranging between *a* and *b*, and a local Gaussian environmental filter with parameters *t*_*opt*_ and *σ*_*opt*_^*2*^, the local trait distribution should follow a truncated Gaussian distribution, such as:

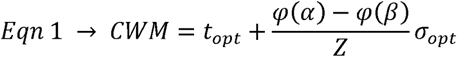

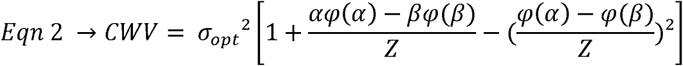

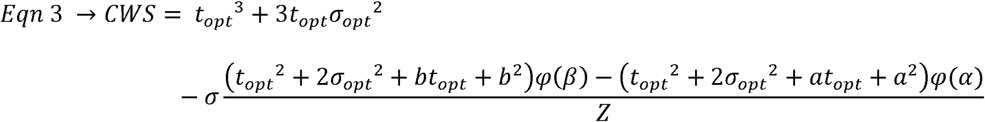

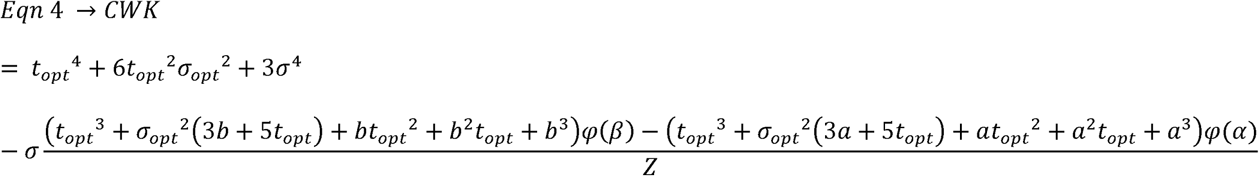

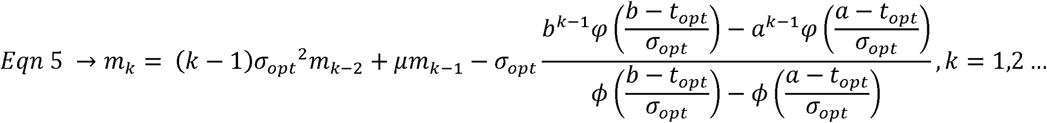

Where 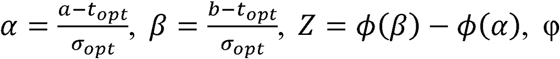 is the probability density function of the standard normal distribution and □ is its cumulative density function (Barr & Sherrill, 1999), *m*_*k*_ is the k^th^ moment of the truncated normal distribution (based on *m*_*-1*_ = 0 and *m*_*0*_ = 1), *a* and *b* are the trait range limits (with *a < b*), *t*_*opt*_ the mean and *σ*_*opt*_ the standard deviation of the Gaussian filtering function, 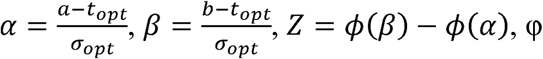 is the probability density function of the standard normal distribution and LJ is its cumulative density function (Barr and Sherrill 1999).

## Appendix S2

R code to simulate and analyze communities.

This example is divided into two parts. The first part simulates communities using the R package *ecolottery* (Munoz et al. 2018), https://cran.r-project.org/web/packages/ecolottery/index.html). In the example of the paper, the parameters used to build communities are the following: species pool with 100.000 individuals belonging to 100 species with equal abundances (i.e., 1000 individuals each), and species trait values drawn from a uniform distribution between *a*=0 and *b*=1. Each community includes 500 individuals and the immigrants establishing in communities are drawn from the species pool. Stabilizing environmental filtering determines establishment probability of immigrants depending on the departure of their trait value *t* from a local optimum *t*_*opt*_. We thus choose a Gaussian filtering function of mean *t*_*opt*_, which varies among communities, and standard deviation *σ*_*opt*_ equal to 0.25. The data frame of simulated communities is called *sim*. The community-weighted mean (CWM) and variance (CWV) are calculated for each community, as well as functional dispersion (Laliberté and Legendre 2010) and Rao’s entropy (Botta-Dukát 2005).

The second section estimates the two parameters *t*_*opt*_ and *σ*_*opt*_ in each community, by comparing observed summary statistics of the community to summary statistics simulated over a broad range of *t*_*opt*_ and *σ*_*opt*_ values, with approximate Bayesian computation (ABC) analysis (*coalesc_abc* function, Munoz et al. 2018).

Finally, the third just presents the code to generate the plot equivalent to the Figure 2 of the manuscript.

~~~
rm(list = ls())
#-----------------------------------------------------------------------------
# PART I/ Generate simulations: uniform species abundances and trait values in
# regional pool, trait values bounded between 0 and 1
#-----------------------------------------------------------------------------
# Installing package from CRAN and loading
install.packages(“ecolottery”) library(ecolottery)
J <- 500
m <- 1
sim <- c()
# Trait range
a <- 0
b <- 1
# Generate a regional pool/metacommunity with equal species abundances and
# uniform trait distribution
# 100000 individuals, 100 species, trait values bounded between 0 and 1
pool <- cbind(1:100000, rep(sample(1:100), 1000), rep(NA, 100000))
colnames(pool) <- c(“ind”, “sp”, “tra”)
t.sp <- runif(1000, min = a, max = b)
pool[, “tra”] <- t.sp[pool[, “sp”]]
# Intraspecific variability
pool[, “tra”] <- rnorm(pool[, “tra”], pool[, “tra”], 0.001)
pool[, “tra”] <- ifelse(pool[, “tra”] > 1, 0.99, pool[, “tra”])
pool[, “tra”] <- ifelse(pool[, “tra”] < 0, 0.01, pool[, “tra”])
# Generate communities with habitat filtering, in 100 communities with
# distinct topt
# sigmaopt is fixed at 0.25
topt <- seq(from = a, to = b, by = (b-a) / 99)
sigmaopt <- 0.25
for(j in 1:length(topt)) {
comm <- coalesc(J, m, pool=pool, traits=NULL, filt = function(x) exp(-(x-topt[j])^2/(2*sigmaopt^2)))
sim <- rbind(sim, cbind(rep(j, nrow(comm$com)), rep(topt[j], nrow(comm$com)), comm$com))
}
# Column names of the metacommunity dataset
colnames(sim) <- c(“com”, “topt”, “ind”, “sp”, “tra”)
# Conversion to data.frame
sim <- as.data.frame(sim)
# Table of species abundances per community
temp <- as.data.frame(table(sim$sp, sim$com))
colnames(temp) <- c(“sp”, “com”, “ab”)
# Relative abundances temp$abrel <- temp$ab / J
# Merging abundances with simulation
library(dplyr)
sim$sp_com <- paste(sim$sp, sim$com, sep=“_”)
temp$sp_com <- paste(temp$sp, temp$com, sep=“_”)
sim <- inner_join(sim, temp[, c(“sp_com”, “ab”, “abrel”)], by=“sp_com”)
# Computing CWM => not weighted by abundances because of intraspecific variability
cwm <- tapply(sim$tra, sim$com, mean)
cwm <- data.frame(“com” = names(cwm), “cwm” = as.numeric(cwm))
sim$com <- as.character(sim$com)
sim <- inner_join(sim, cwm, by = “com”)
# Computing CWV => not weighted by abundances because of intraspecific variability
cwv <- tapply(sim$tra, sim$com, var)
cwv <- data.frame(“com” = names(cwv), “cwv” = as.numeric(cwv))
sim <- inner_join(sim, cwv, by = “com”)
# Computing Fdis and Rao
library(FD)
FD_com <- c()
for(i in 1:length(unique(sim$com))){
com <- sim[which(sim$com == unique(sim$com)[i]), c(“ind”, “tra”, “abrel”)]
com <- com[!duplicated(com),]
tra_ind <- com[, c(“ind”, “tra”)]
rownames(tra_ind) <- tra_ind$ind
tra_ind <- tra_ind[, “tra”, drop = FALSE]
ab_ind <- rep(1, nrow(com))
names(ab_ind) <- com$ind
tmp <- dbFD(tra_ind, a = ab_ind, w.abun = FALSE)
FD_com <- rbind(FD_com, c(unique(sim$com)[i], as.numeric(tmp$FDis), as.numeric(tmp$RaoQ)))
}
FD_com <- data.frame(FD_com)
colnames(FD_com) <- c(“com”, “fdis”, “rao”)
sim <- inner_join(sim, FD_com, by = “com”)
sim$fdis <- as.numeric(as.character(sim$fdis))
sim$rao <- as.numeric(as.character(sim$rao))
# Plot showing correlations between CWV and functional diversity metrics
library(GGally)
ggpairs(sim[!duplicated(sim$com), c(“cwv”, “fdis”, “rao”)]) +
theme_classic()
#-----------------------------------------
# PART II/ ABC-based parameter estimation
#-----------------------------------------
require(vegan)
# Function to compute 6 summary statistics: the four first orders of community
# weighted moments (mean, variance, skewness and kurtosis) and two taxonomic
# statistics (specific richness and Shannon diversity index)
f.sumstats <- function(com){
array(dimnames = list(c(“cwm”, “cwv”, “cws”, “cwk”, “S”, “Es”)),
c(mean(com[, 3]), var(com[, 3]), e1071::skewness(com[, 3]),
e1071::kurtosis(com[, 3]), vegan::specnumber(table(com[, 2])),
vegan::diversity(table(com[, 2]))))
}
# Observed summary statistics
ss.obs <- c()
for(i in 1:length(unique(sim$com))) {
comm <- sim[which(sim$com == unique(sim$com)[i]), c(“ind”, “sp”, “tra”)]
ss.obs[[i]] <- f.sumstats(comm)
}
comm.sd <- unlist(lapply(ss.obs, function(x) sqrt(x[“cwv”])))
comm.cwm <- unlist(lapply(ss.obs, function(x) x[“cwm”]))
# Possibility to reconstruct the pool of species from communities composition
# true_pool <- pool
# pool <- sim[, c(“ind”, “sp”, “tra”)]
# Filtering function
filt_gaussian <- function(t, params) exp(-(t-params[1])^2/(2*params[2]^2))
# Parameters values
params <- data.frame(rbind(c(min(pool[, “tra”]), max(pool[, “tra”])),
c(min(comm.sd), sd(pool[, “tra”]))))
row.names(params) <- c(“topt”, “sigmaopt”)
# Number of values to sample in prior distributions
nb.samp <- 10^6
# ABC estimation of the parameters based on summary statistics of the observed
# community
# The function makes vary the migration rate, m, and the parameters of
# environmental filtering defined in params
res <- c()
for(i in 1:length(unique(sim$com))) {
comm <- sim[which(sim$com == unique(sim$com)[i]), c(“ind”, “sp”, “tra”)]
res[[i]] <- coalesc_abc(comm, pool, f.sumstats = f.sumstats,
filt.abc = filt_gaussian,
params=params, nb.samp = 1000, parallel = TRUE,
tol = 1, pkg = c(“e1071”,“vegan”),
method = “neuralnet”)
}
# Mean estimated values of the parameters
topt.abc <- unlist(lapply(res, function(x) weighted.mean(x$abc$adj.value[, “topt”], w = x$abc$weights)))
sigmaopt.abc <- unlist(lapply(res, function(x) weighted.mean(x$abc$adj.value[, “sigmaopt”], w = x$abc$weights)))
m.abc <- unlist(lapply(res, function(x) weighted.mean(x$abc$adj.value[, “m”], w = x$abc$weights)))
# Adding topt and sigmaopt to original dataset
library(dplyr)
topt_sim <- data.frame(“com” = unique(sim$com), “topt_abc” = topt.abc)
sim <- inner_join(sim, topt_sim, by = “com”)
sigmaopt_sim <- data.frame(“com” = unique(sim$com), “sigmaopt_abc” = sigmaopt.abc)
sim <- inner_join(sim, sigmaopt_sim, by = “com”)
m_sim <- data.frame(“com” = unique(sim$com), “m_abc” = m.abc)
sim <- inner_join(sim, m_sim, by = “com”)
##----------------------------------------------------------------------
# PART III/ Plots
##----------------------------------------------------------------------
simplot <- sim[!duplicated(sim$com),]
# Slope tests
mobs <- lm(cwm ~ topt, data = simplot)
mabc <- lm(topt_abc ~ topt, data = simplot)
# Quadratic regression for CWV
mvobs <- lm(cwv ~ topt + I(topt^2), data = simplot)
mvabc <- lm(sigmaopt_abc ~ topt + I(topt^2), data = simplot)
obs_slope <- paste0(“Slope coefficient = ”,round(summary(mobs)$coefficients[2, 1], 2), “ [”, round(confint(mobs)[2, 1], 2), ”, ”,round(confint(mobs)[2, 2], 2), “]”)
abc_slope <- paste0(“Slope coefficient = ”,round(summary(mabc)$coefficients[2, 1], 2), “ [”, round(confint(mabc)[2, 1], 2), ”, ”,round(confint(mabc)[2, 2], 2), “]”)
# Four panels
par(mfrow = c(2, 2), mai = c(1, 0.9, 0.1, 0.3))
y_lim <- c(round((min(simplot$cwm, simplot$topt_abc)*2/2), 2), round((max(simplot$cwm, simplot$topt_abc)*2/2), 2))
# CWM ~ topt
plot(simplot$topt, simplot$cwm, xlab = “”, ylab = “”, xaxt = “n”, yaxt = “n”,
xlim = c(0, 1), ylim = c(y_lim[1], y_lim[2]),
col = “firebrick3”, pch = 16, cex = 1.5)
axis(1, cex.axis = 1.4)
mtext(expression(“t”[“opt”]), side = 1, line = 2.2, cex = 2)
axis(2, cex.axis = 1.4)
mtext(“CWM”, side = 2, line = 2.2, cex = 2)
abline(0, 1, lwd = 1)
legend(x = -0.1, y = 11/10 * y_lim[2],
bty = “n”, legend = “a”, cex = 2, col = “black”)
legend(−0.2, 10/10 * y_lim[2], obs_slope, bty = “n”, cex = 2)
# topt_ABC ~ topt
plot(simplot$topt, simplot$topt_abc, xlab = “”, ylab = “”, xaxt = “n”,
yaxt = “n”, xlim = c(0, 1), ylim = c(y_lim[1], y_lim[2]),
col = “dodgerblue”, pch = 16, cex = 1.5)
axis(1, cex.axis = 1.4)
mtext(expression(“t”[“opt”]), side = 1, line = 2.2, cex = 2)
axis(2, cex.axis = 1.4)
mtext(expression(hat(“t”)[“opt”]), side = 2, line = 2.2, cex = 2)
abline(0, 1, lwd = 1)
legend(x = -0.1, y = 11/10 * y_lim[2],
bty = “n”, legend = “b”, cex = 2, col = “black”)
legend(−0.2, 10/10 * y_lim[2], abc_slope, bty = “n”, cex = 2)
# CWV ~ topt
y_lim <- c(round((min(simplot$cwv, simplot$sigmaopt_abc^2) * 9/10), 2),
round((max(simplot$cwv, simplot$sigmaopt_abc^2) * 11/10), 2))
cwv_diff <- mean(abs(simplot$cwv - 0.25^2))
cwv_diff <- scales::scientific_format()(cwv_diff)
plot(simplot$topt, simplot$cwv, xlab = “”, ylab = “”, xaxt = “n”, yaxt = “n”,
xlim = c(0, 1), ylim = c(y_lim[1], y_lim[2]),
col = “firebrick3”, pch = 16, cex = 1.5)
legend(x = -0.1, y = y_lim[2] * 11/10, bty = “n”, legend = “c”, cex = 2, col = “black”)
axis(1, cex.axis = 1.4)
mtext(expression(“t”[“opt”]), side = 1, line = 2.2, cex = 2)
axis(2, cex.axis = 1.4)
mtext(“CWV”, side = 2, line = 2.2, cex = 2)
abline(0.25^2, 0, cex = 2)
# sigmaopt_ABC^2 ~ topt
sigmaopt_abc_diff <- mean(abs(simplot$sigmaopt_abc^2 - 0.25^2))
sigmaopt_abc_diff <- scales::scientific_format()(sigmaopt_abc_diff)
plot(simplot$topt, simplot$sigmaopt_abc^2, xlab = “”, ylab = “”, xaxt = “n”,
yaxt = “n”, xlim = c(0, 1), ylim = c(y_lim[1], y_lim[2]),
col = “dodgerblue”, pch = 16, cex = 1.5)
legend(x = -0.1, y = y_lim[2] * 11/10,
bty = “n”, legend = “d”, cex = 2, col = “black”)
axis(1, cex.axis = 1.4)
mtext(expression(“t”[“opt”]), side = 1, line = 2.2, cex = 2)
axis(2, cex.axis = 1.4)
mtext(expression(hat(sigma)[“opt”]^“2”), side = 2, line = 2.2, cex = 2)
abline(0.25^2, 0, cex = 2)
~~~

## Appendix S3.

Variation of CWV values with CWM (top, red color), and of 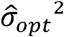 with *t*_*opt*_ and (bottom, blue color), for two sets of simulated communities with constant (left, *σ*_*opt*_ = 0.25) or varying (*σ*_*opt*_ from 0.25 to 0.05, peaking at *t*_*opt*_ = 0.5) intensity of environmental filtering.

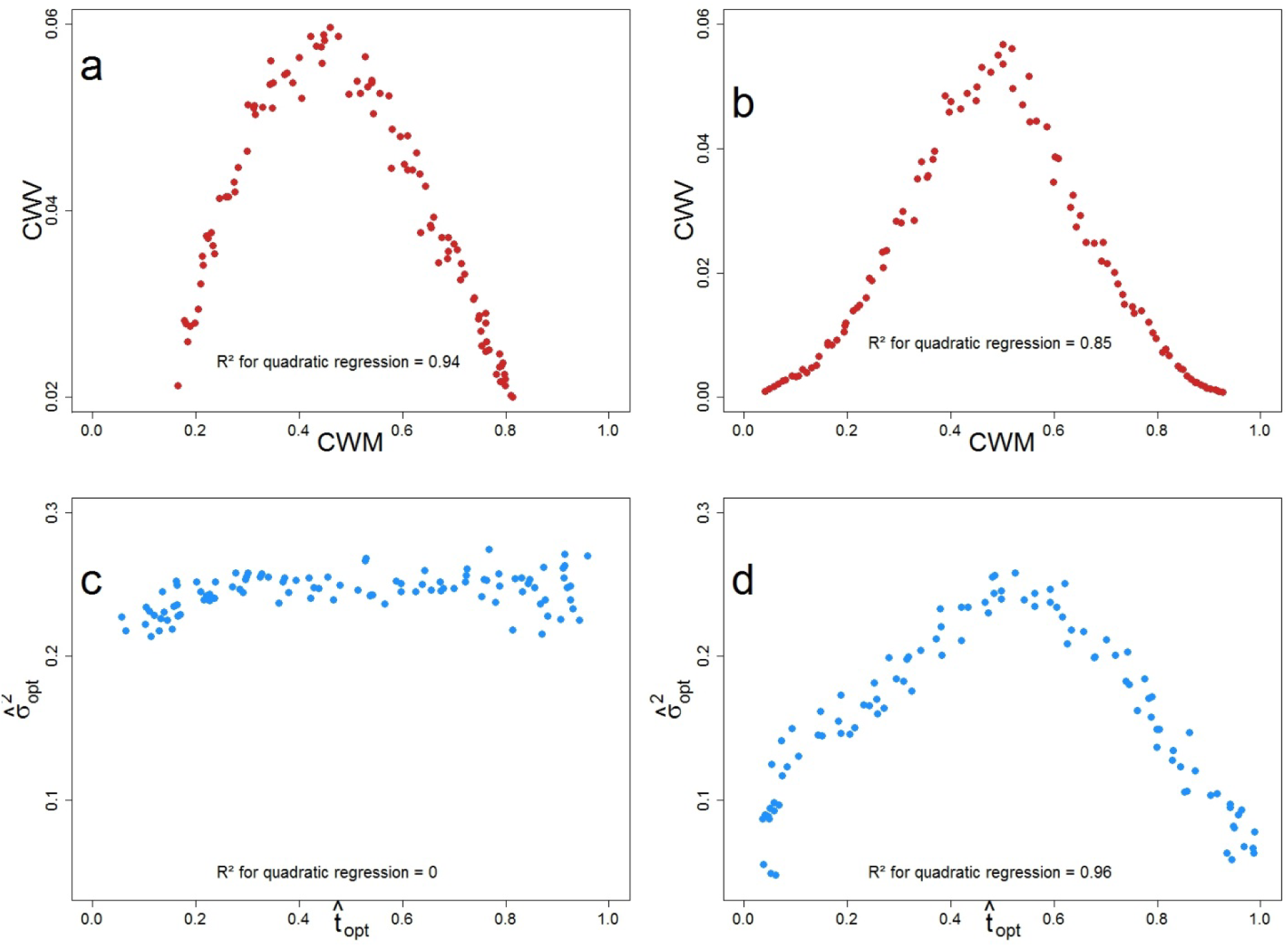

## Appendix S4.

Analysis of simulated communities with constant and strong environmental filtering (*σ*_*opt*_ = 0.05), low intraspecific variability (*σ* = 0.001 for each species trait value) and uniform distribution of species pool abundances.

The left red curves show the variation of CWM (top) and CWV (bottom) according to *t*_*opt*_. The right blue curves show the estimated 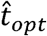(top) and 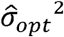 (bottom) values according to *t*_*opt*_. The black solid line represents equality of CWM and CWV to the parameters of environmental filtering (*t*_*opt*_ and *σ*_*opt*_, respectively). Slope coefficients and the associated confidence intervals of the linear regression equations between CWM/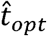 and *t*_*opt*_ are displayed in panel (a) and (b). The mean of the difference between *σ*_*opt*_^*2*^ and CWV (c) is comparable to the difference between *σ*_*opt*_^*2*^ and 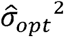 (d) (respectively 6.02e-2 and 6e-2).

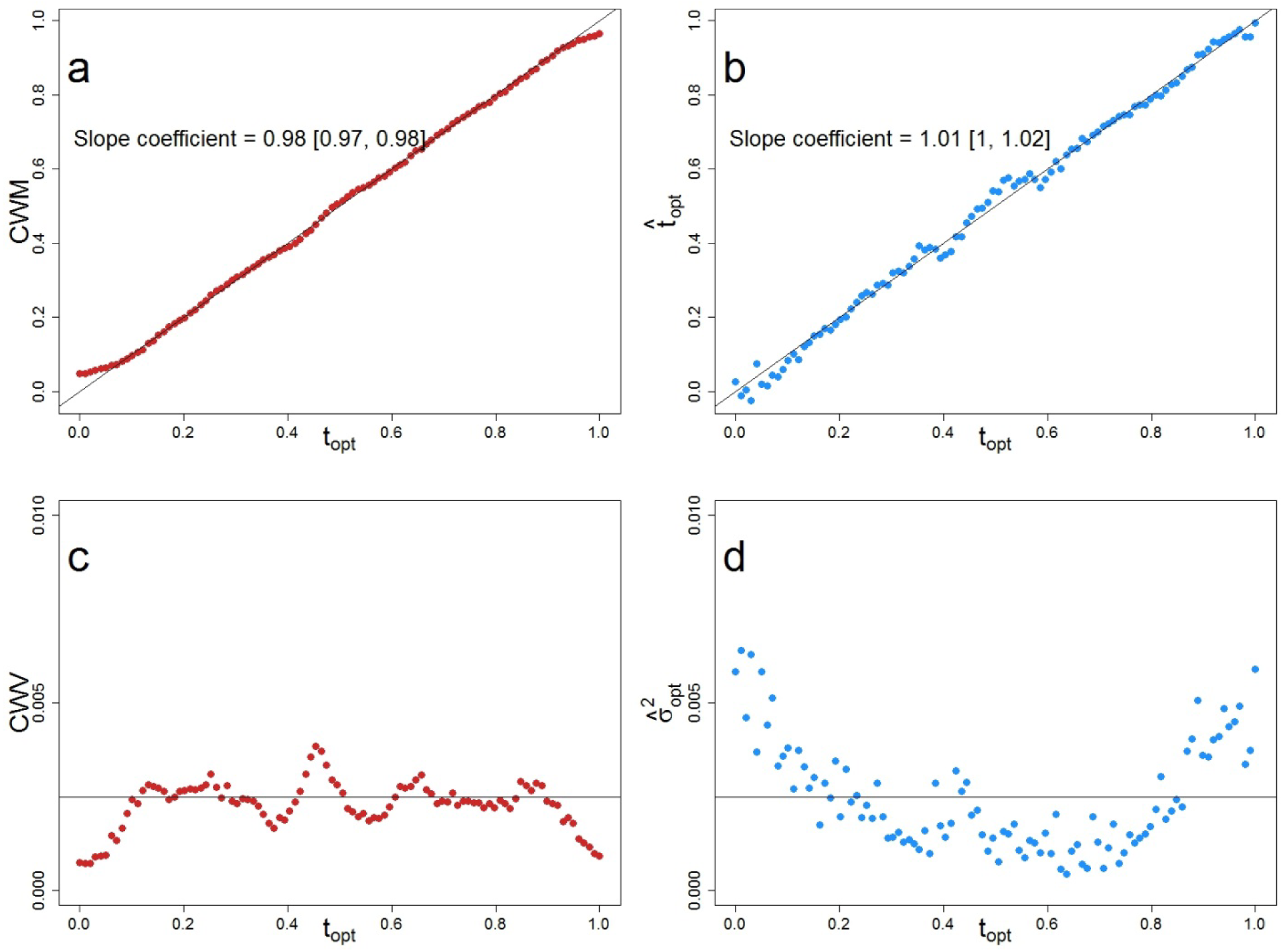

## Appendix S5.

Community-weighted skewness (CWS) and kurtosis (CWK) of simulated communities. CWS is calculated for community *j* as

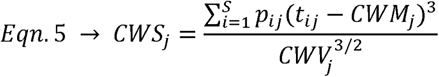

and CWK is calculated for community *j* as

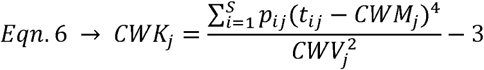

Where *S* is the number of species in community *j*, *p*_*ij*_ is the relative abundance of species *i* in community *j*, *t*_*ij*_ is the average trait value of species *i* in community *j*.

Panel (a) displays the variation of CWS and panel (b) of CWK in simulated communities according to *t*_*opt*_, with uniform species pool abundances and constant environmental filtering (σ_opt_ = 0.25; same dataset as in Fig. 2).

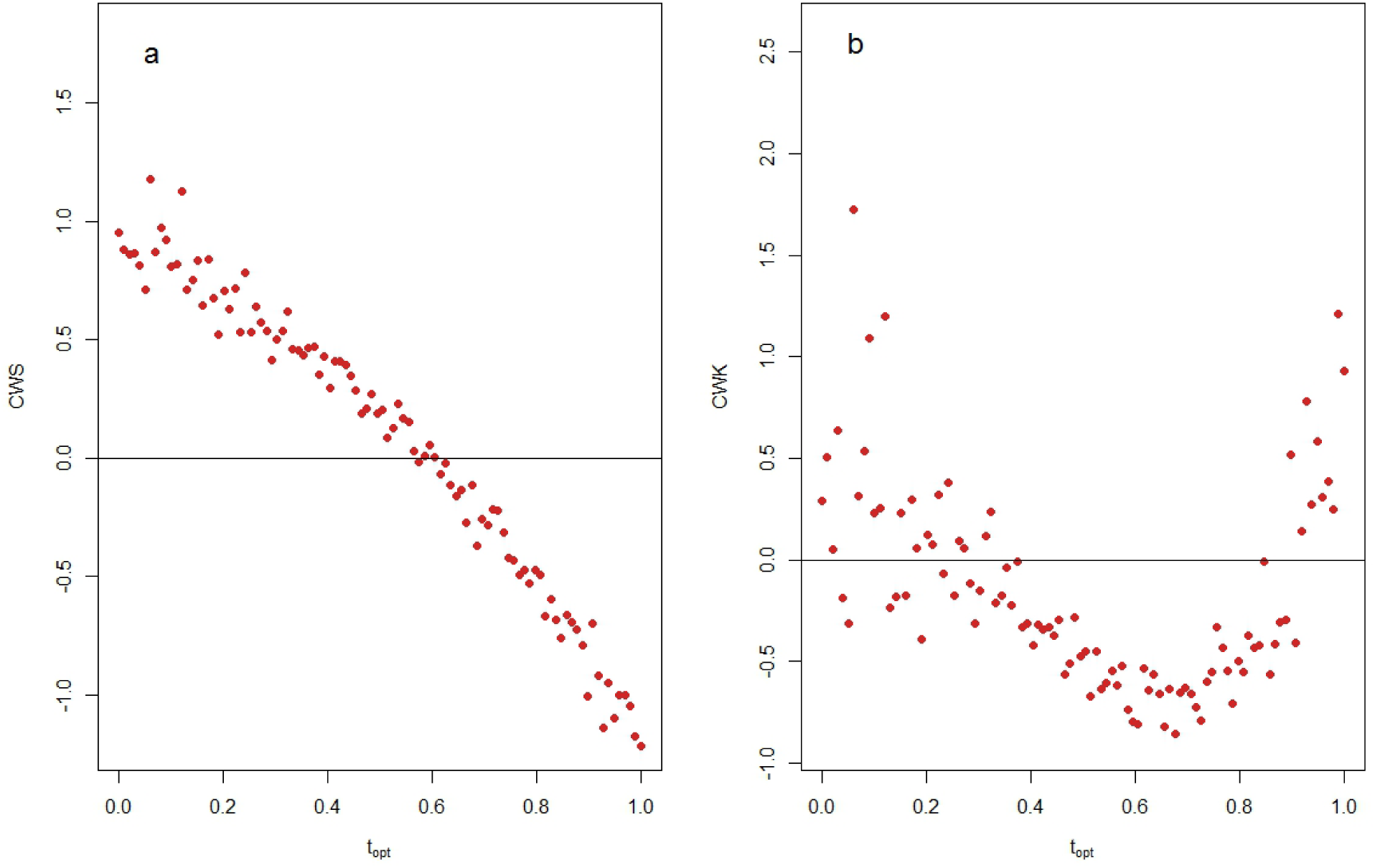

## Appendix S6.

Variation of CWM and CWV according to *t*_*opt*_, with constant environmental filtering intensity (*σ*_*opt*_ = 0.25), large intraspecific variability (*σ*_*opt*_ = 0.1 for each species trait value) and a uniform distribution of species pool abundances.

The left part (red curves) shows the variation of CWM (top) and CWV (bottom) according to *t*_*opt*_. The right part (blue curves) shows the estimated topt 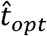 (top) and 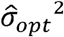 (bottom) values. The black solid line represents equality of CWM and CWV to the parameters of environmental filtering (*t*_*opt*_ and *σ*_*opt*_, respectively). Slope coefficients and the associated confidence intervals of the linear regression equations between CWM/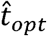 and *t*_*opt*_ are displayed in panel (a) and (b). The mean of the difference between *σ*_*opt*_^*2*^ and CWV (c) is twice higher than the difference between *σ*_*opt*_^*2*^ and 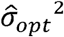 (d) (respectively 2.3e-2 and 4.78e-3).

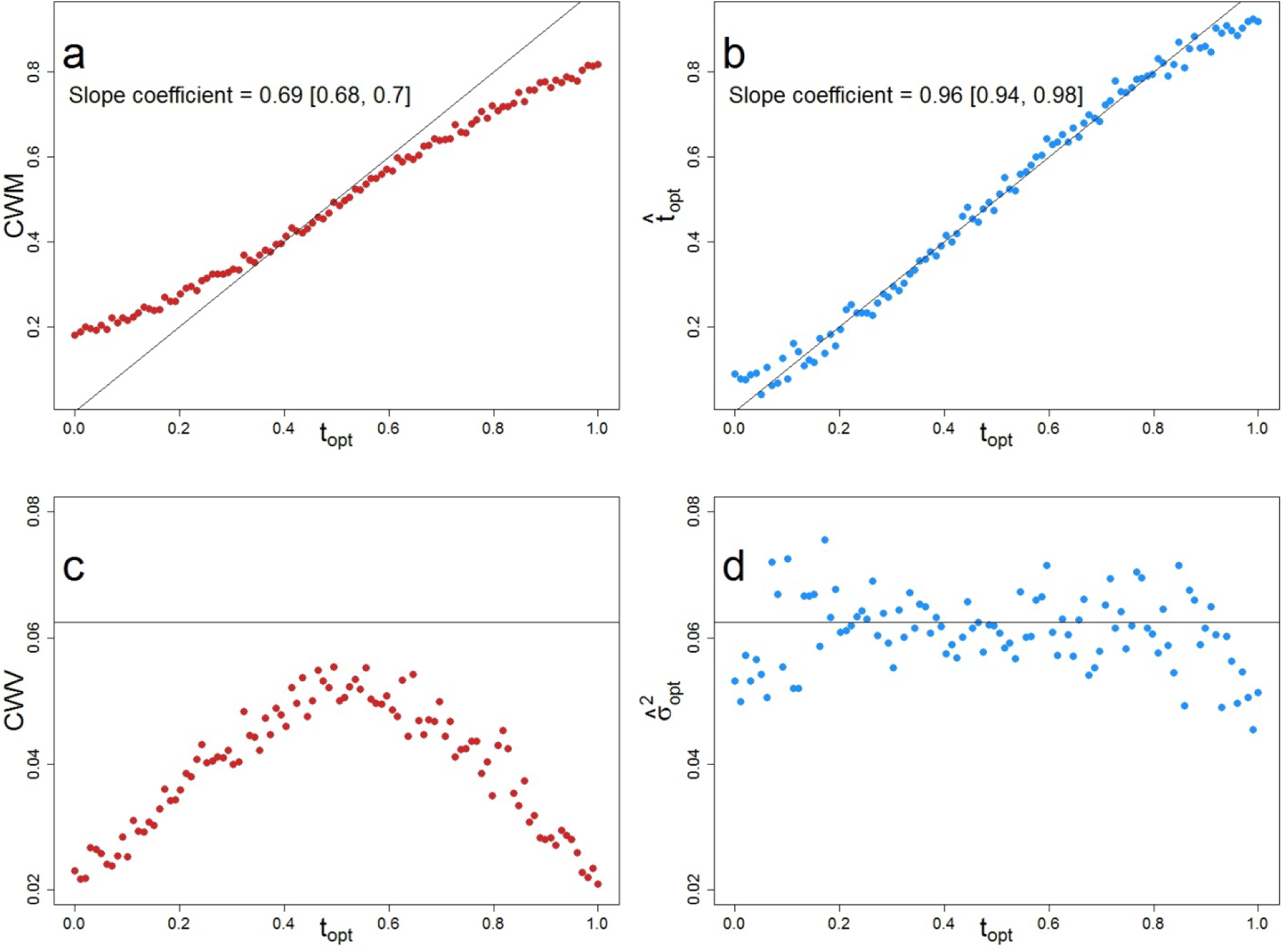

## Appendix S7.

Variation of CWM and CWV according to *t*_*opt*_, with constant environmental filtering intensity (σ_opt_ = 0.25) and a log-series (biodiversity parameter θ = 50) distribution of species pool abundances.

The left red curves show the variation of CWM (top) and CWV (bottom) according to *t*_*opt*_. The right blue curves show the estimated topt 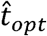 (top) and 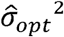 (bottom) values. The black solid line represents equality of CWM and CWV to the parameters of environmental filtering (*t*_*opt*_ and σ_opt_^2^, respectively). Slope coefficients and the associated confidence intervals of the linear regression equations between CWM / 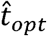 and *t*_*opt*_ are displayed in panel (a) and (b). The mean of the difference between *σ*_*opt*_^*2*^ and CWV (c) is twice higher than the difference between *σ*_*opt*_^*2*^ and 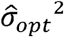 (d) (respectively 2.18e-2 and 4.79e-3).

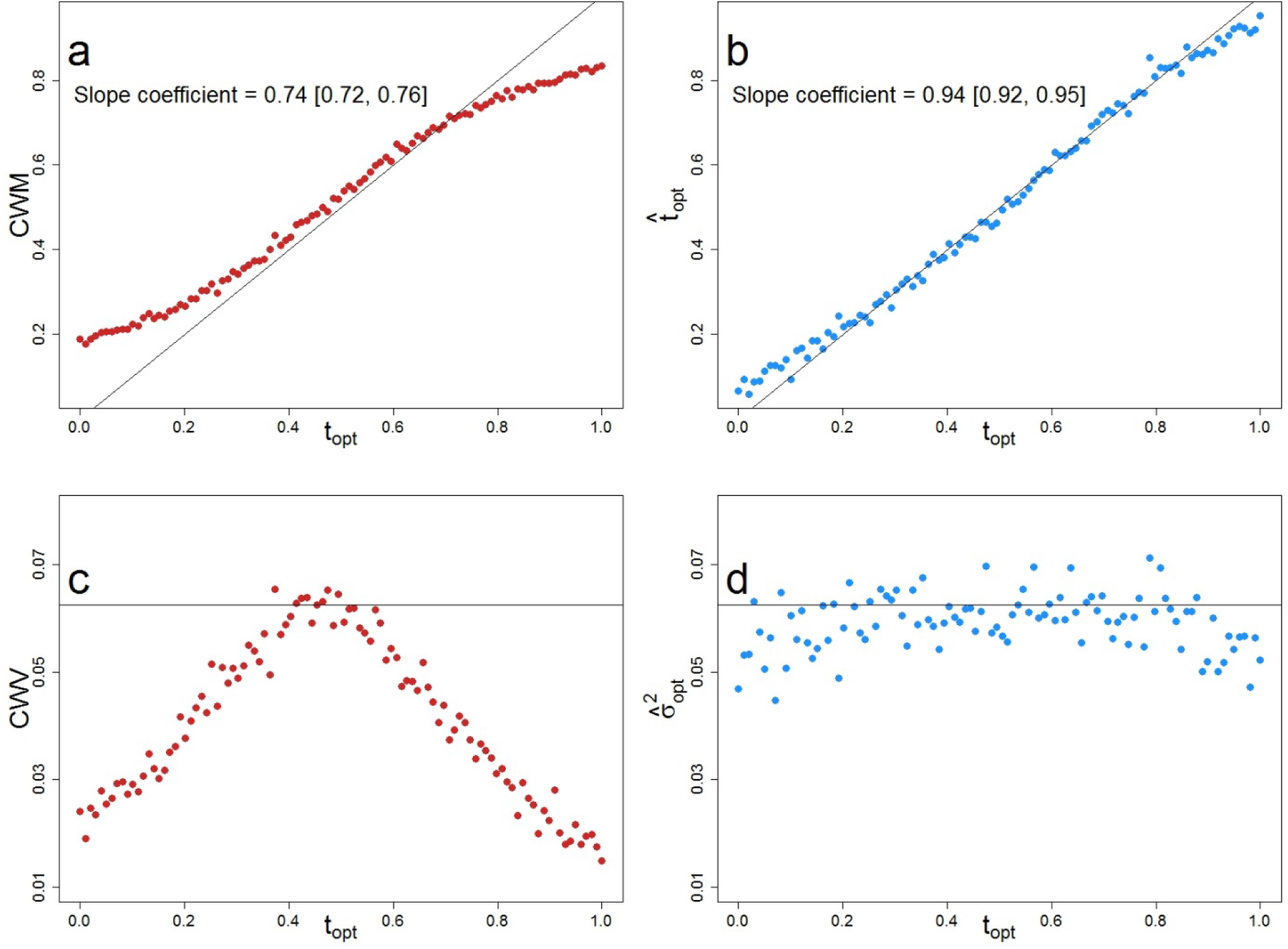

## Appendix S8.

Influence of environmental filtering intensity and trait range extent on the departure of CWM from *t*_*opt*_.

The global deviation of CWM from *t*_*opt*_ over the trait range (average distance between CWM and 1:1 line), summarizes the influence of the trait range limits on CWM over the whole environmental gradient. Blue, resp. black, points represent simulations with a trait range on [0; 1], resp. [0; 2], and varying filtering intensity (*σ*_*opt*_, on abscissa).

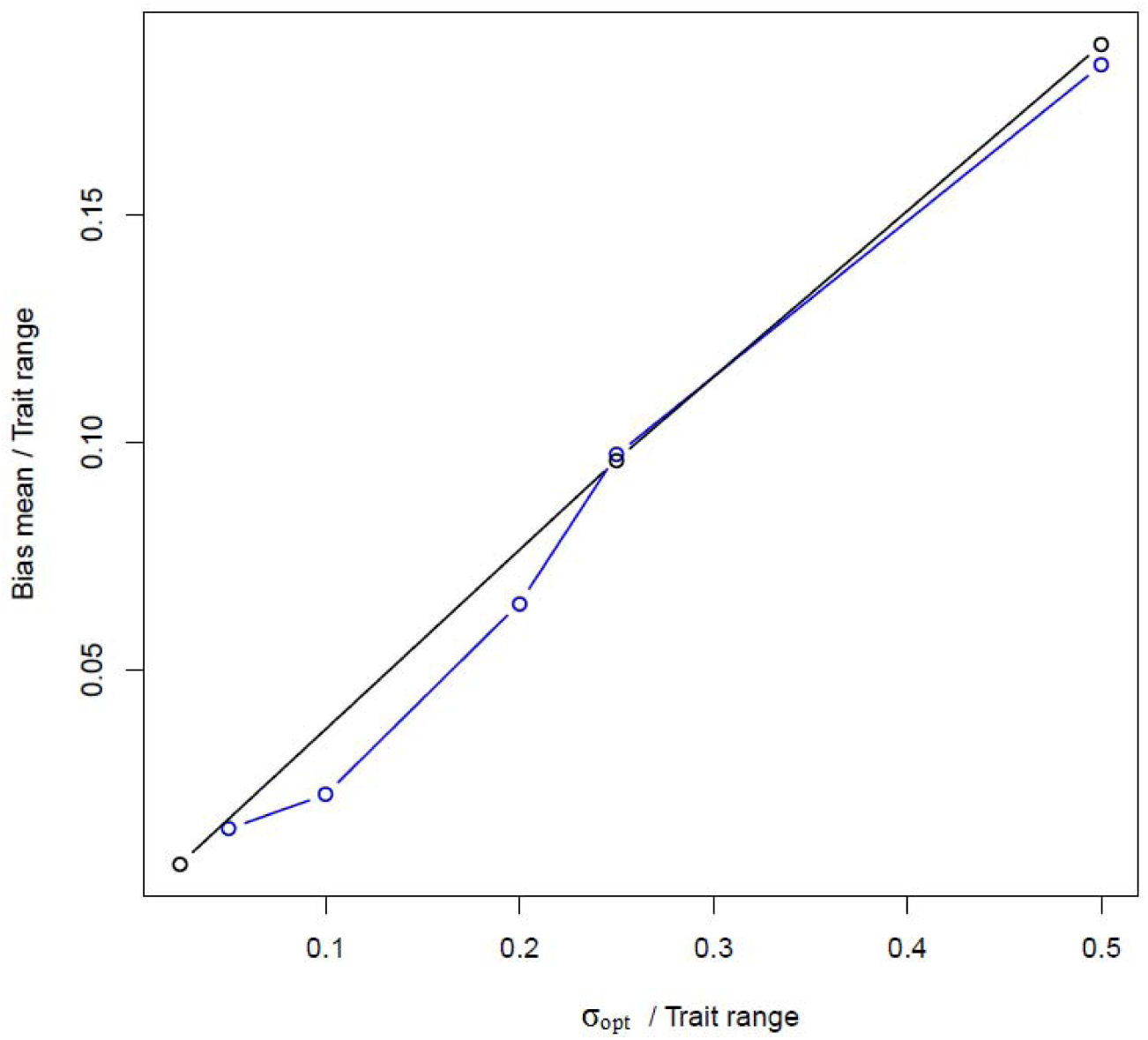

## Appendix S9.

Observed weighted skewness in communities (CWS) for the *aravo* dataset. Panels a & b display the result for the alpine plant communities for N_mass_ (panel a) and SLA (panel b).

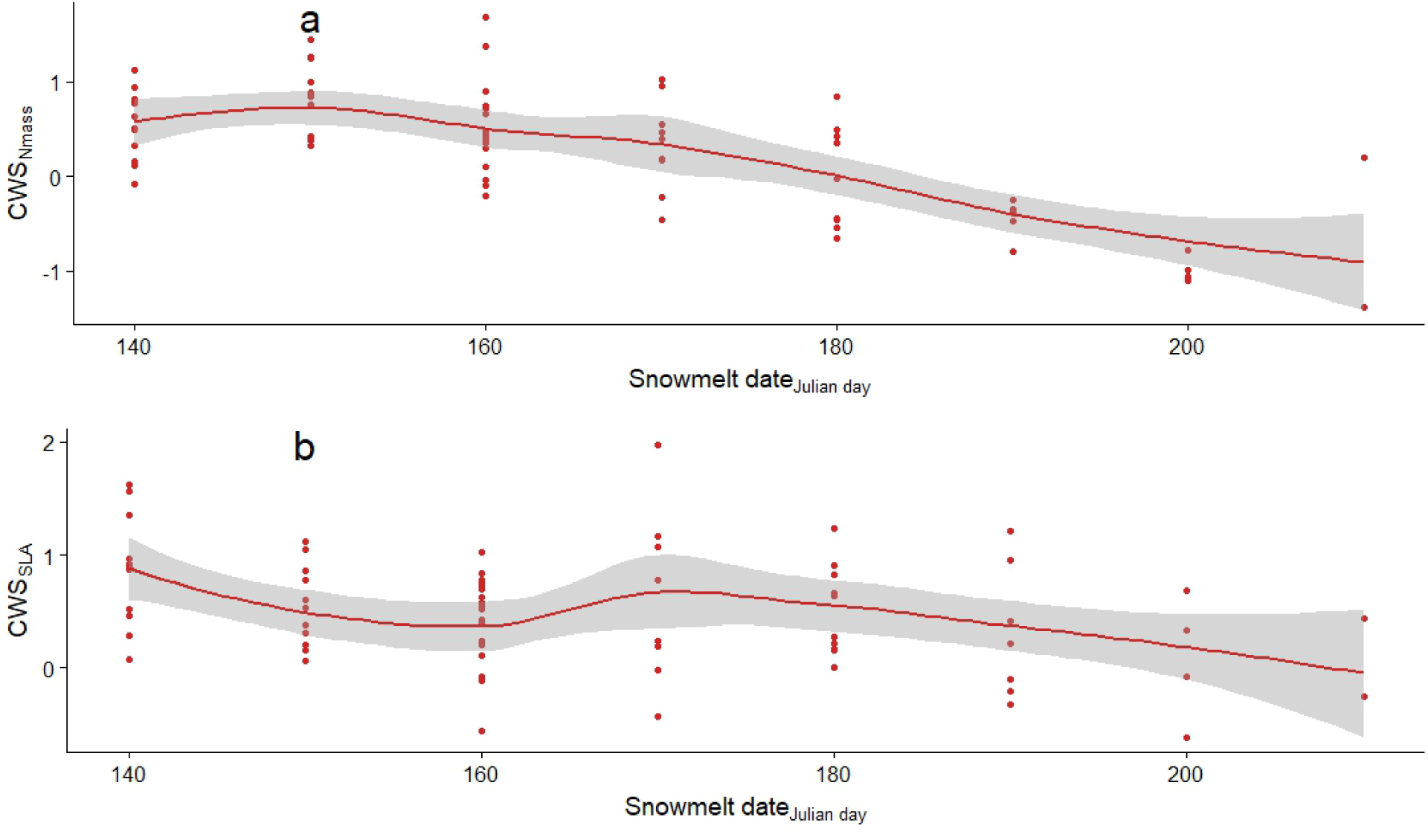

## Appendix S10.

Variation of functional dispersion, Rao’s quadratic entropy and 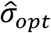 according to *t*_*opt*_, with constant environmental filtering intensity (σ_opt_ = 0.25) and auniform distribution of species pool abundances.

The first two red curves show the variation of functional dispersion (first panel) and Rao’s quadratic entropy (second panel) according to *t*_*opt*_. The third right blue curve shows the estimated 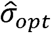 values. The black solid line represents equality of CWV to the parameter of environmental filtering σ_opt_,. The fraction of variance explained by quadratic regression between the three metrics and σ_opt_ (R^2^) are displayed.

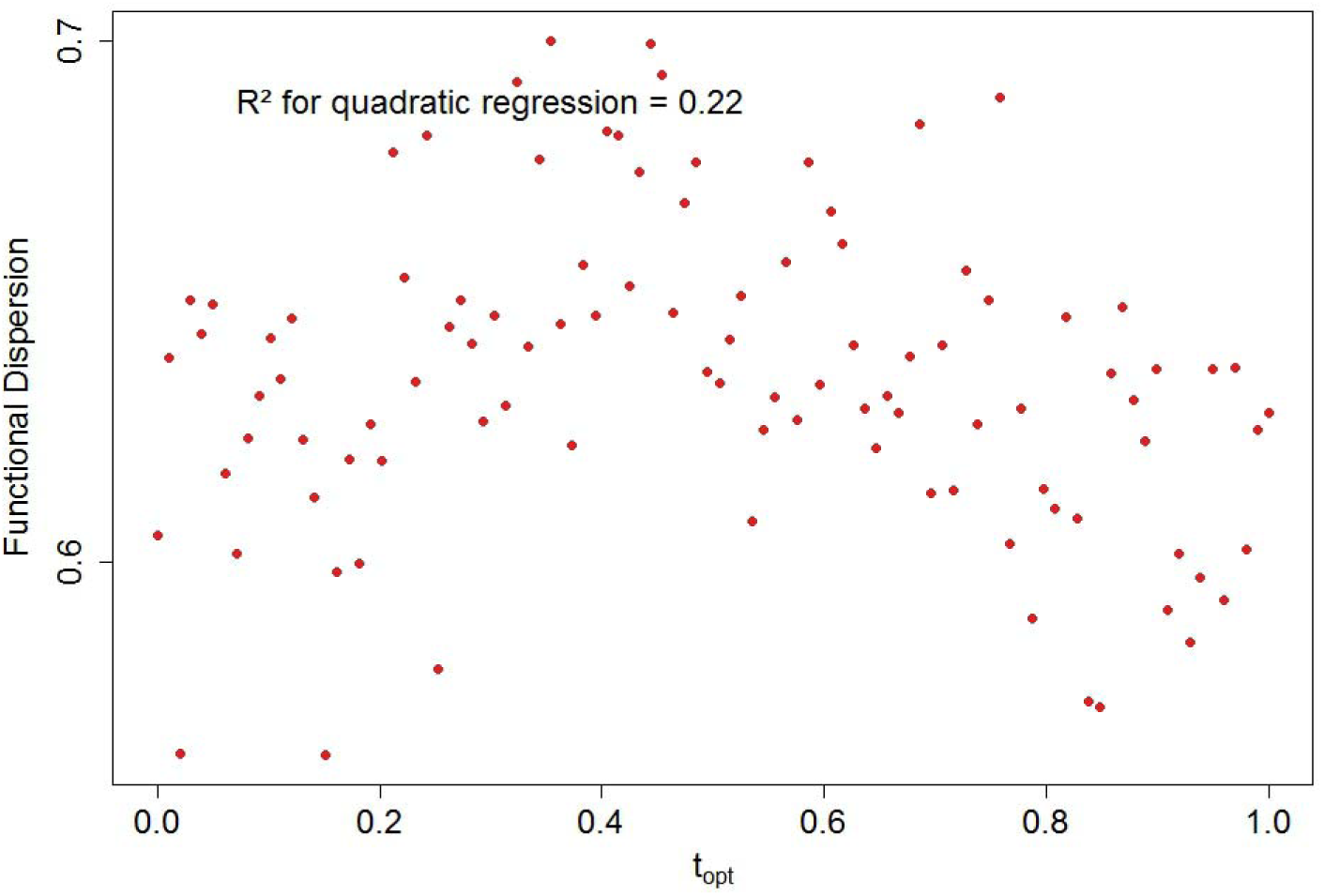
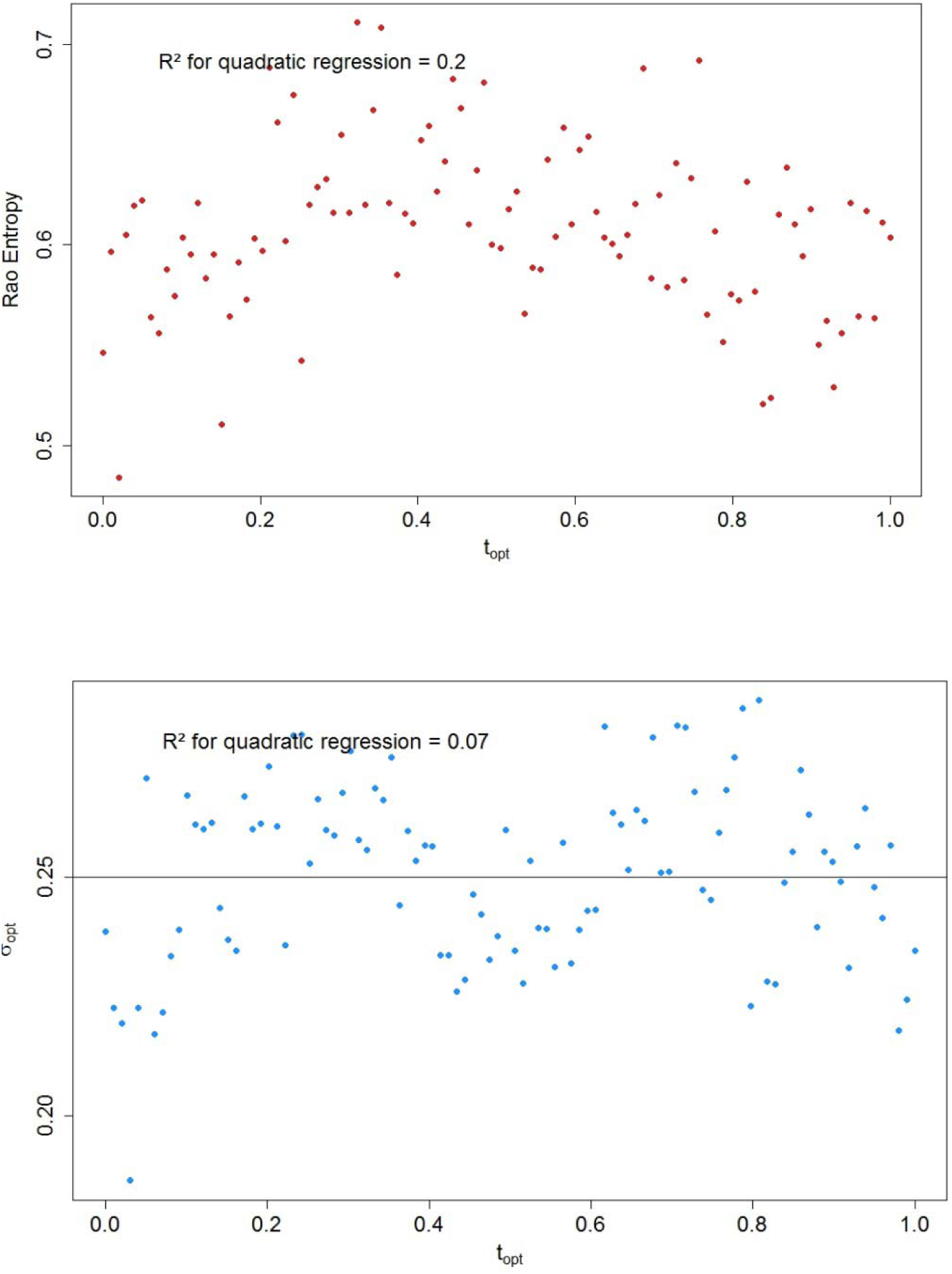

## Appendix S11.

Variation in CWM and CWV values (left, red color), and of estimated 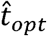 and 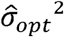 (right, blue color), for simulated communities along *t*_*opt*_ gradient with the observed species pool.

Communities were simulated with constant environmental filtering (*σ*_*opt*_ = 0.25), uniform distribution of trait values and uniform abundances in the species pool. Top figures (a) and (b) represent CWM and topt, and figures (c) and (d) represent CWV and 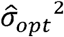. The 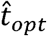 and 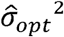 values were obtained with the ABC approach and correctly estimated the *t*_*opt*_ and *σ*_*opt*_^*2*^ values (b and d). The species pool used for ABC estimation of the parameters corresponds to the actual sum of observed communities. Conversely, CWM departed from *t*_*opt*_ and CWV was below *σ*_*opt*_^*2*^ when the influence of the trait range limits increased at the extremes. The black solid line represents equality of CWM and CWV to the parameters of environmental filtering (*t*_*opt*_ and *σ*_*opt*_^*2*^, respectively). Slope coefficients and the associated confidence intervals of the linear regression equations between CWM / 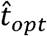 and *t*_*opt*_ are displayed in panel (a) and (b). The mean of the difference between *σ*_*opt*_^*2*^ and CWV (c) is twice higher than the difference between *σ*_*opt*_^*2*^ and 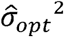 (respectively 2.23e-2 and 8.91e-3).

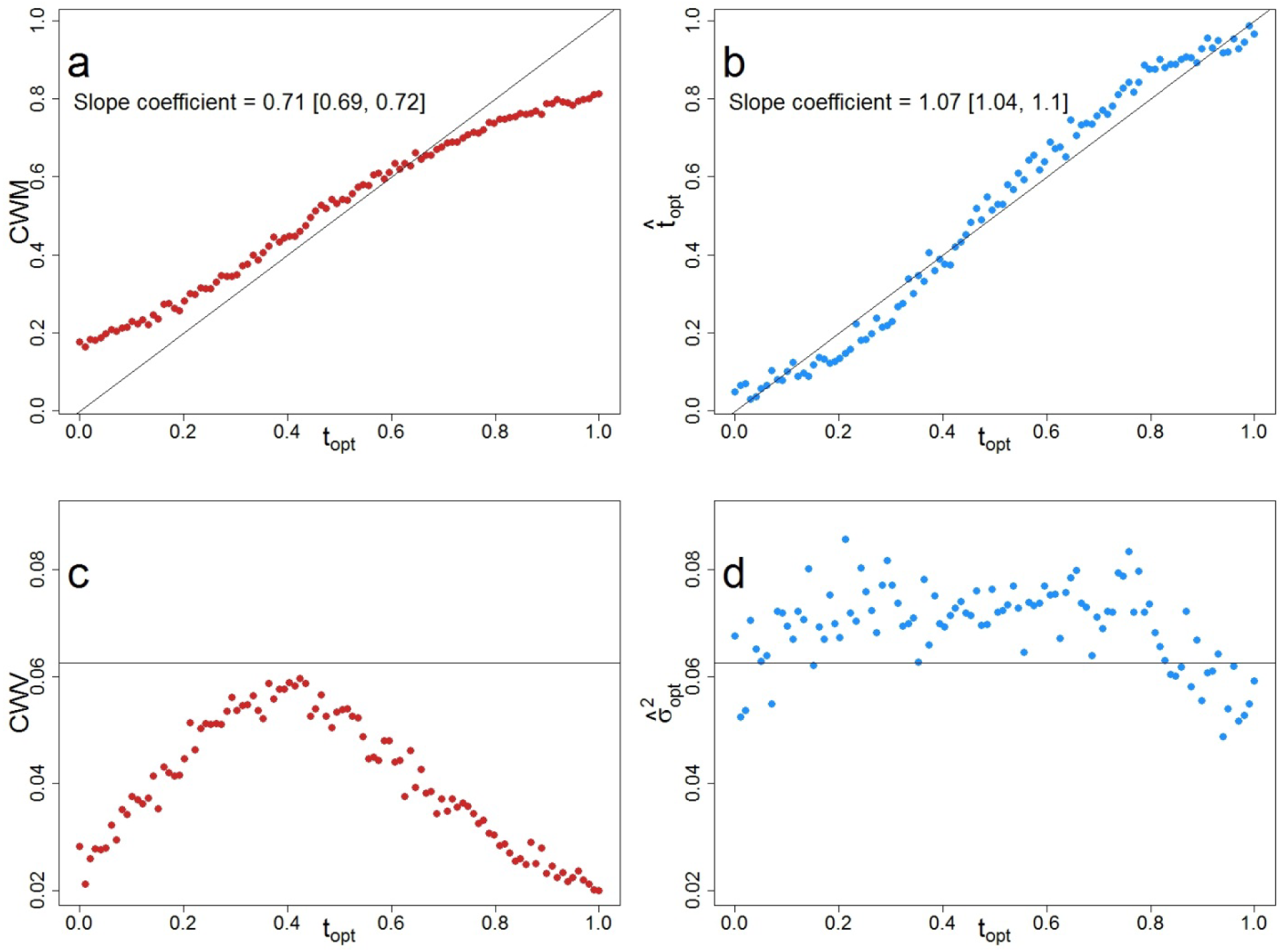

## References

Ackerly, D. D. and Cornwell, W. K. 2007. A trait-based approach to community assembly: partitioning of species trait values into within- and among-community components. - Ecol. Lett. 10: 135–145.

Adler, P. B. et al. 2013. Trait-based tests of coexistence mechanisms. - Ecol. Lett. 16: 1294–1306.

Alpert, P. 2005. The Limits and Frontiers of Desiccation-Tolerant Life1. - Integr. Comp. Biol. 45: 685–695.

Bernard-Verdier, M. et al. 2012. Community assembly along a soil depth gradient: contrasting patterns of plant trait convergence and divergence in a Mediterranean rangeland (H Cornelissen, Ed.). - J. Ecol. 100: 1422–1433.

Borgy, B. et al. 2017a. Sensitivity of community-level trait–environment relationships to data representativeness: A test for functional biogeography. - Glob. Ecol. Biogeogr. 26: 729–739.

Borgy, B. et al. 2017b. Plant community structure and nitrogen inputs modulate the climate signal on leaf traits. - Glob. Ecol. Biogeogr. 26: 1138–1152.

Botta-Dukát, Z. 2005. Rao’s quadratic entropy as a measure of functional diversity based on multiple traits. - J. Veg. Sci. 16: 533–540.

Callaway, R. M. et al. 2002. Positive interactions among alpine plants increase with stress. - Nature 417: 844–848.

Chesson, P. 2000. Mechanisms of maintenance of species diversity. - Annu. Rev. Ecol. Syst.: 343–366.

Choler, P. 2005. Consistent Shifts in Alpine Plant Traits along a Mesotopographical Gradient. - Arct. Antarct. Alp. Res. 37: 444–453.

Cingolani, A. M. et al. 2007. Filtering processes in the assembly of plant communities: Are species presence and abundance driven by the same traits? - J. Veg. Sci. 18: 911–920.

Colwell, R. K. and Lees, D. C. 2000. The mid-domain effect: geometric constraints on the geography of species richness. - Trends Ecol. Evol. 15: 70–76.

Colwell, R. K. et al. 2004. The Mid-Domain Effect and Species Richness Patterns:What Have We Learned So Far? - Am. Nat. 163: E1–E23.

Cornwell, W. K. et al. 2006. A trait-based test for habitat filtering: Convex hull volume. - Ecology 87: 1465–1471.

Csilléry, K. et al. 2010. Approximate Bayesian computation (ABC) in practice. - Trends Ecol. Evol. 25: 410–418.

Dias, A. T. C. et al. 2013. An experimental framework to identify community functional components driving ecosystem processes and services delivery (S Lavorel, Ed.). - J. Ecol. 101: 29–37.

Ehrlén, J. and Eriksson, O. 2000. Dispersal limitation and patch occupancy in forest herbs. - Ecology 81: 1667–1674.

Enquist, B. J. et al. 2015. Chapter Nine-Scaling from Traits to Ecosystems: Developing a General Trait Driver Theory via Integrating Trait-Based and Metabolic Scaling Theories. - Adv. Ecol. Res. 52: 249–318.

Fierer, N. et al. 2014. Seeing the forest for the genes: using metagenomics to infer the aggregated traits of microbial communities. - Front. Microbiol. 5:614.

Forrestel, E. J. et al. 2017. Different clades and traits yield similar grassland functional responses. - Proc. Natl. Acad. Sci. 114: 705–710.

Garnier, E. et al. 2016. Plant functional diversity: organism traits, community structure, and ecosystem properties. - Oxford University Press.

Gross, N. et al. 2017. Functional trait diversity maximizes ecosystem multifunctionality. - Nat. Ecol. Evol. 1: 0132.

Hubbell, S. P. 2001. The unified neutral theory of biodiversity and biogeography Princeton University Press Princeton.

Jiménez-Alfaro, B. et al. 2018. History and environment shape species pools and community diversity in European beech forests. - Nat. Ecol. Evol.: 1.

Koch, G. W. et al. 2004. The limits to tree height. - Nature 428: 851.

Kraft, N. J. B. et al. 2015. Community assembly, coexistence and the environmental filtering metaphor (J Fox, Ed.). - Funct. Ecol. 29: 592–599.

Laliberté, E. and Legendre, P. 2010. A distance-based framework for measuring functional diversity from multiple traits. - Ecology 91: 299–305.

Lamanna, C. et al. 2014. Functional trait space and the latitudinal diversity gradient. - Proc. Natl. Acad. Sci. 111: 13745–13750.

Laughlin, D. C. et al. 2018. Survival rates indicate that correlations between community-weighted mean traits and environments can be unreliable estimates of the adaptive value of traits. - Ecol. Lett.21: 411-421.

Leibold, M. A. et al. 2004. The metacommunity concept: a framework for multi-scale community ecology: The metacommunity concept. - Ecol. Lett. 7: 601–613.

Lepš, J. et al. 2011. Community trait response to environment: disentangling species turnover vs intraspecific trait variability effects. - Ecography 34: 856–863.

Lessard, J.-P. et al. 2011. Strong influence of regional species pools on continent-wide structuring of local communities. - Proc. R. Soc. Lond. B Biol. Sci.: rspb20110552.

Lessard, J.-P. et al. 2016. Process-based species pools reveal the hidden signature of biotic interactions amid the influence of temperature filtering. - Am. Nat. 187: 75.

Letten, A. D. et al. 2013. The mid-domain effect: it’s not just about space (R Pearson, Ed.). - J. Biogeogr. 40: 2017–2019.

Levine, J. M. and Murrell, D. J. 2003. The community-level consequences of seed dispersal patterns. - Annu. Rev. Ecol. Evol. Syst. 34: 549–574.

Lomolino, M. V. et al. 2006. Biogeography. - Sinauer Associates Sunderland, MA.

Loranger, J. et al. 2018. What makes trait–abundance relationships when both environmental filtering and stochastic neutral dynamics are at play? - Oikos in press.

Mayfield, M. M. and Levine, J. M. 2010. Opposing effects of competitive exclusion on the phylogenetic structure of communities: Phylogeny and coexistence. - Ecol. Lett. 13: 1085–1093.

Mcgill, B. et al. 2006. Rebuilding community ecology from functional traits. - Trends Ecol. Evol. 21: 178–185.

Mitchell, R. M. et al. 2017. Species’ traits do not converge on optimum values in preferred habitats. - Oecologia: 1–11.

Mittelbach, G. G. and Schemske, D. W. 2015. Ecological and evolutionary perspectives on community assembly. - Trends Ecol. Evol. 30: 241–247.

Munoz, F. et al. 2018. ecolottery: Simulating and assessing community assembly with environmental filtering and neutral dynamics in R. - Methods Ecol. Evol. 9: 693–703.

Muscarella, R. and Uriarte, M. 2016. Do community-weighted mean functional traits reflect optimal strategies? - Proc. R. Soc. B Biol. Sci. 283: 20152434.

Newbold, T. et al. 2012. Mapping Functional Traits: Comparing Abundance and Presence-Absence Estimates at Large Spatial Scales (F de Bello, Ed.). - PLoS ONE 7: e44019.

Pärtel, M. et al. 2011. Dark diversity: shedding light on absent species. - Trends Ecol. Evol. 26: 124–128.

Patrick, C. J. and Brown, B. L. 2018. Species Pool Functional Diversity Plays a Hidden Role in Generating β- Diversity. - Am. Nat. 191: E159–E170.

Pey, B. et al. 2014. Current use of and future needs for soil invertebrate functional traits in community ecology. - Basic Appl. Ecol. 15: 194–206.

Ricotta, C. and Moretti, M. 2011. CWM and Rao’s quadratic diversity: a unified framework for functional ecology. - Oecologia 167: 181–188.

Shipley, B. 2010. From plant traits to vegetation structure: chance and selection in the assembly of ecological communities. - Cambridge University Press.

Siefert, A. et al. 2015. A global meta-analysis of the relative extent of intraspecific trait variation in plant communities. - Ecol. Lett. 18: 1406–1419.

Spasojevic, M. J. et al. 2018. Integrating species traits into species pools. - Ecology 99: 1265–1276.

Umaña, M. N. et al. 2015. Commonness, rarity, and intraspecific variation in traits and performance in tropical tree seedlings (K Suding, Ed.). - Ecol. Lett.18: 1329-1337.

Violle, C. et al. 2007. Let the concept of trait be functional! - Oikos 116: 882–892.

Violle, C. et al. 2012. The return of the variance: intraspecific variability in community ecology. - Trends Ecol. Evol. 27: 244–252.

Violle, C. et al. 2014. The emergence and promise of functional biogeography. - Proc. Natl. Acad. Sci. 111: 13690–13696.

Violle, C. et al. 2015. Vegetation ecology meets ecosystem science: Permanent grasslands as a functional biogeography case study. - Sci. Total Environ. 534: 43–51.

Violle, C. et al. 2017. Functional Rarity: The Ecology of Outliers. - Trends Ecol. Evol. 32:356-367.

Webb, C. T. et al. 2010. A structured and dynamic framework to advance traits-based theory and prediction in ecology. - Ecol. Lett. 13: 267–283.

Weiher, E. et al. 1998. Community assembly rules, morphological dispersion, and the coexistence of plant species. - Oikos: 309–322.

Weiher, E. et al. 2011. Advances, challenges and a developing synthesis of ecological community assembly theory. - Philos. Trans. R. Soc. B Biol. Sci. 366: 2403–2413.

Zobel, M. 1997. The relative of species pools in determining plant species richness: an alternative explanation of species coexistence? - Trends Ecol. Evol. 12: 266–269.

## Reference

Barr, D. R. and Sherrill, E. T. 1999. Mean and variance of truncated normal distributions. - Am. Stat. 53: 357–361.

